# A Cre-Driver Rat Model for Anatomical and Functional Analysis of Glucagon *(Gcg)*-Expressing Cells in the Brain and Periphery

**DOI:** 10.1101/2022.10.10.511573

**Authors:** Huiyuan Zheng, Lorena López-Ferreras, Jean-Phillipe Krieger, Stephen Fasul, Valentina Cea Salazar, Natalia Valderrama Pena, Karolina P. Skibicka, Linda Rinaman

## Abstract

**Objective:** The glucagon gene (*Gcg*) encodes preproglucagon, which is cleaved to form glucagon-like peptide 1 (GLP1) and other mature signaling molecules implicated in metabolic functions. To date there are no transgenic rat models available for precise manipulation of GLP1-expressing cells in the brain and periphery.

**Methods:** To visualize and manipulate *Gcg*-expressing cells in rats, CRISPR/Cas9 was used to express *iCre* under control of the *Gcg* promoter. Gcg-Cre rats were bred with tdTomato reporter rats to tag *Gcg*-expressing cells. Cre-dependent AAVs and RNAscope *in situ* hybridization were used to evaluate the specificity of *iCre* expression by GLP1 neurons in the caudal nucleus of the solitary tract (cNTS) and intermediate reticular nucleus (IRt), and by intestinal and pancreatic secretory cells. Food intake was assessed in heterozygous (Het) Gcg-Cre rats after chemogenetic stimulation of cNTS GLP1 neurons expressing an excitatory DREADD.

**Results:** While genotype has minimal effect on body weight or composition in chow-fed Gcg-Cre rats, homozygous (Homo) rats have lower plasma glucose levels. In neonatal and adult Gcg-Cre/tdTom rats, reporter-labeled cells are present in the cNTS and IRt, and in additional brain regions (e.g., basolateral amygdala, piriform cortex) that lack detectable *Gcg* mRNA in adults but display transient developmental or persistently low *Gcg* expression. Compared to wildtype (WT) rats, hindbrain *Gcg* mRNA and GLP1 protein in brain and plasma are markedly reduced in Homo Gcg-Cre rats. Chemogenetic stimulation of cNTS GLP1 neurons reduced overnight chow intake in males but not females, the effect in males was blocked by antagonism of central GLP1 receptors, and hypophagia was enhanced when combined with a subthreshold dose of cholecystokinin-8 to stimulate gastrointestinal vagal afferents.

**Conclusions:** Gcg-Cre rats are a novel and valuable experimental tool for analyzing the development, anatomy, and function of *Gcg*-expressing cells in the brain and periphery. In addition, Homo Gcg-Cre rats are a unique model for assessing the role of *Gcg*-encoded proteins in glucose homeostasis and energy metabolism.

**Highlights:**

- A transgenic Gcg-Cre rat model expresses *iCre* under control of the *Gcg* promoter
- *iCre* is expressed by GLP1-positive cells in the hindbrain, pancreas, and intestine
- Additional brain regions display transient and/or very low levels of *Gcg* expression
- +/+ Gcg-Cre rats display a marked knockdown of *Gcg* mRNA and GLP1 protein
- Chemogenetic activation of *Gcg* neurons suppresses food intake in +/- Gcg-Cre rats

## 1.0 Introduction

Preproglucagon (PPG) is a precursor protein encoded by the glucagon gene *(Gcg)*. PPG is cleaved in a tissue-specific manner to form several mature peptides, including glucagon, glucagon-like peptide 1 (GLP1), GLP2, oxyntomodulin, glicentin, and glicentin-related pancreatic polypeptide [1]. In humans, rats, and mice, GLP1 and other *Gcg*-encoded peptides are released from pancreatic islet alpha cells and intestinal enteroendocrine L-cells to regulate insulin secretion, glucose metabolism, and gastrointestinal functions [2][3]. The GLP1 receptor-mediated actions of these peripherally-released peptides form the basis for highly effective pharmacotherapies to treat type 2 diabetes [4].

In the adult rodent brain, *Gcg* is expressed by a subset of olfactory bulb interneurons and by glutamatergic hindbrain neurons whose cell bodies occupy the caudal nucleus of the solitary tract (cNTS) and intermediate reticular nucleus (IRt) [5][6][7]. *Gcg*-expressing hindbrain neurons are GLP1-immunopositive, are sensitive to metabolic state, project to multiple subcortical regions of the central nervous system (CNS), and are implicated in neuroendocrine, autonomic, and behavioral responses to cognitive and physical stressors [8][9][10][11][12][13][14]. *Gcg*-expressing and/or GLP1-positive cells in the pancreas, intestine, and hindbrain are similarly distributed in rodents and primates, including humans [15][16][1][17], and a recent genomic characterization of *Gcg* and related receptor genes suggests a high degree of functional conservation across mammalian species [18].

Current knowledge regarding the structure and function of *Gcg*-expressing cells is based largely on preclinical research using rodent models, with especially valuable contributions arising from experiments using transgenic mice to visualize and manipulate *Gcg*-expressing cells with a high degree of anatomical and cellular specificity [19][20][21][22][23][24][25]. In some cases, research using rodent models is more translationally valid if rats are used instead of or in addition to mice [26][27]. Rats are genetically and metabolically more similar to humans than are mice, and their larger size makes them a preferred model for experimental procedures that use surgical approaches to target the circulatory system, CNS, and visceral organs. Rats also have a richer behavioral repertoire and are less stress-responsive than mice, making them generally more tractable for behavioral and metabolic research. Thus, the availability of a transgenic rat model for targeted analysis and manipulation of *Gcg*-expressing cells in the brain and periphery will advance research in systems neuroscience, digestive physiology, energy metabolism, and behavior.

Towards this goal, we used the CRISPR/Cas9 technique [28] to generate a novel Sprague Dawley (SD) knock-in rat line, Gcg-Cre, in which cells express *iCre* recombinase under control of the *Gcg* promoter. We also used a cross-breeding strategy to generate Gcg-Cre/tdTom reporter rats to facilitate identification and spatial analysis of *Gcg* expression within the neonatal and adult brain, intestine, and pancreas. Here we report data characterizing and validating this unique transgenic rat model, including evidence for transient developmental *Gcg* expression in brain regions that lack detectable *Gcg* expression in adult rodents. We further report that hindbrain *Gcg* mRNA and GLP1 protein levels in brain and plasma are markedly reduced in homozygous (Homo) Gcg-Cre rats. In heterozygous (Het) rats, GLP1 protein levels are moderately reduced in plasma but not in brain. Finally, we report results demonstrating that chemogenetic excitation of cNTS GLP1 neurons inhibits chow intake in male but not female Het Gcg-Cre rats in a central GLP1 receptor (GLP1R)-mediated manner, and enhances the hypophagic effect of a behaviorally subthreshold dose of cholecystokinin-8 (CCK).

## 2.0 Materials and methods

Animal work was approved by the Institutional Animal Care and Use Committees of Florida State University and the Sahlgrenska Academy at University of Gothenburg, Sweden, and was performed in adherence with the Guide for the Care and Use of Laboratory Animals [29]. Unless otherwise noted, adult rats were pair-housed in tub cages with wood-chip bedding in a vivarium with a 12:12 hour light/dark cycle (lights off from 1700-0500h) and *ad libitum* access to rodent chow (Purina 5001) and water.

### 2.1 Generation of Gcg-Cre knockin rats

The rat *Gcg* gene (glucagon, Gene ID: 24952) includes one transcript. To avoid disrupting *Gcg* expression, *iCre* was inserted after the final coding sequence of exon 6 before the 3′ UTR. Contracted work to generate the model was performed by Biocytogen Co., Ltd (Beijing, China), using previously described CRISPR/Cas9 technology [28]. An internal ribosome entry site (IRES) for *iCre* expression was used to permit endogenous promoter-driven cellular expression of *Gcg*, with a lower level of *iCre* expression. The targeting strategy is schematized in **Figure 1A**. Single guide (sg)RNA plasmids were constructed and confirmed by DNA sequencing. One-cell zygotes were microinjected with Cas9 mRNA, sgRNA, and targeting vectors, and then transferred into pseudopregnant SD rats. Four pups were positively confirmed as F0 founder chimera rats by PCR product sequencing. After reaching sexual maturity, one female F0 rat was bred with a wildtype (WT) male SD rat to obtain a litter of germline transmission heterozygous (Het; +/-) F1 pups. The correct targeting of *iCre* in nine F1 pups was confirmed using southern blot (**Fig. 1B,C**) and PCR.

**Figure 1.**
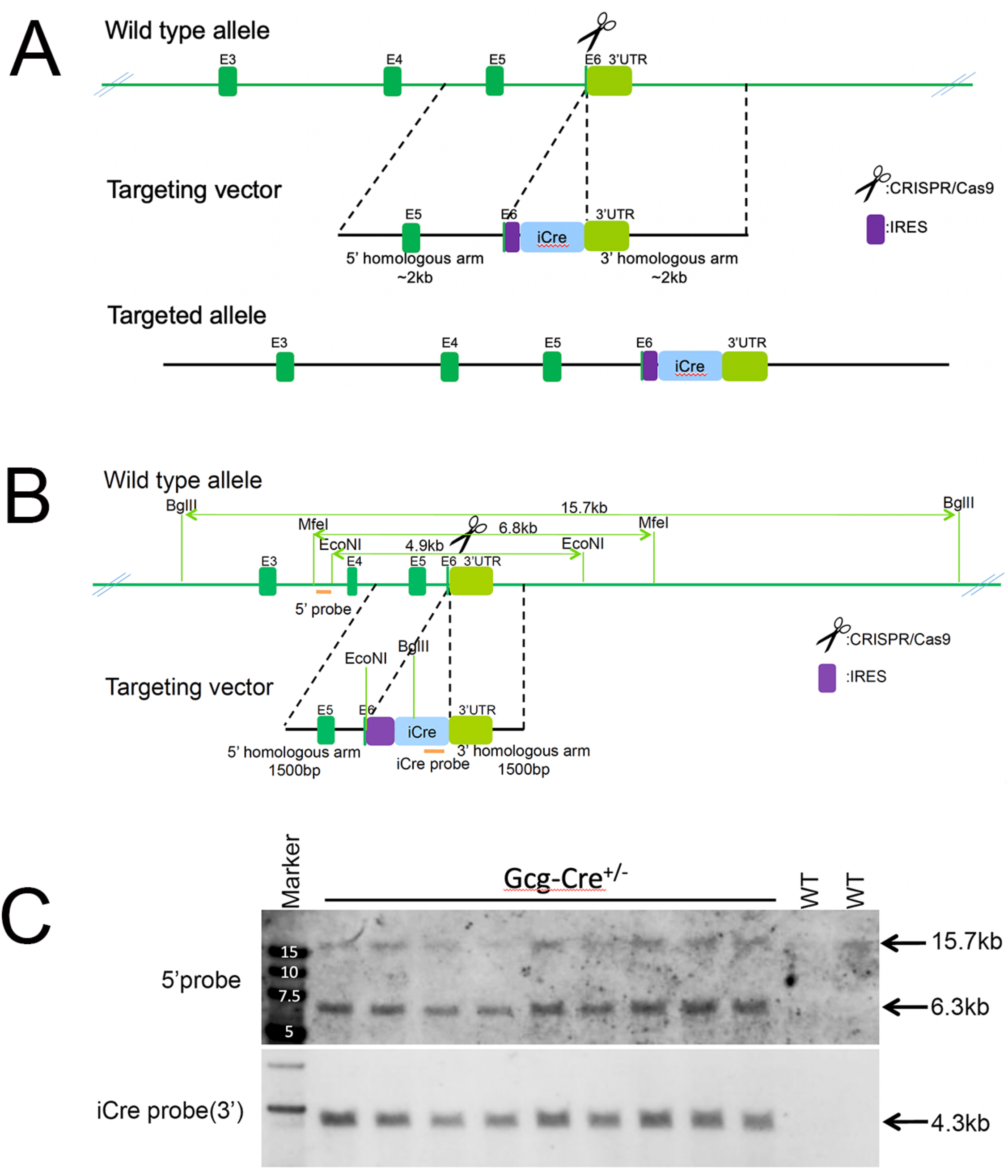
**A**, Targeting strategy for creating Gcg-Cre rats. **B**, Southern blot restriction enzyme strategy and confirmation (**C**) in nine F1 Gcg-Cre^+/-^ rats. Genomic DNA digestion via BgIII, 5’ probe yields two potential DNA fragments (6.3kb targeted allele, 15.7kb WT allele). DNA digestion via EcoNI, iCre probe (3’) yields one DNA fragment (4.3kb targeted allele, no band for WT allele).

### 2.2 Breeding colony establishment and maintenance

#### 2.2.1 FSU Colony

Two F1 Het Gcg-Cre rats (one male, one female) shipped from China to Florida State University (FSU) were used to establish a local breeding colony, with offspring genotypes determined from weanling rat ear punches, using two PCR primer sets (**Table 1)**. To maintain the FSU colony on an outbred SD genetic background, non-sibling Het Gcg-Cre rats are bred together to yield homozygous (Homo, +/+), Het (+/-), and WT (-/-) progeny, with backcrossing of male and female Homo Gcg-Cre rats to WT Gcg-Cre and outbred SD rats (Envigo, Indianapolis, IN) to generate litters of Het rat pups.

**Table 1.**
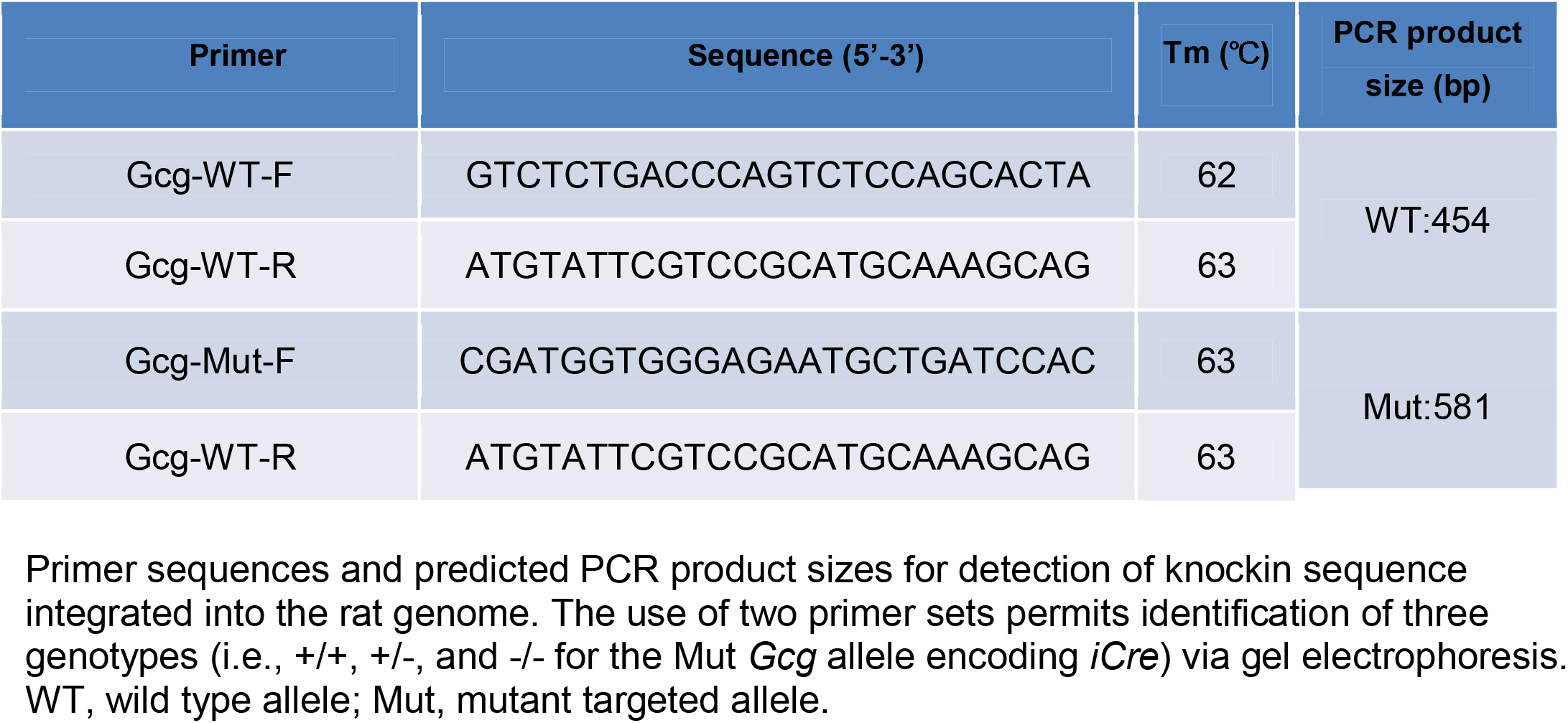
Primer sequences and predicted PCR product sizes.

#### 2.2.2 Janvier Colony

Het Gcg-Cre male rats were shipped from China to the Janvier facility in France to rederive the line. Rats were genotyped on arrival and biopsies stored. Gcg-Cre males were naturally mated with Janvier SD females. Resulting embryos were collected and successively washed with sterile medium. Washed embryos were then transferred to pseudopregnant SD females, which were housed in isolators while their health was monitored. Resulting offspring genotype was determined from tail biopsies by Transnetyx, Inc. (Cordova, TN) using Real Time (RT)-PCR. Some WT and Het progeny born at Janvier were shipped to the Univ. of Gothenburg and used there for behavioral experiments. Other Het Gcg-Cre rats were bred at Janvier, and their genotyped offspring used for body weight measurements. Other WT, Het, and Homo Gcg-Cre rats were shipped from Janvier to the Univ. of Gothenburg and used there for qRT-PCR analysis of *Gcg* and *iCre* gene expression.

### 2.3 Body weight, body composition, and plasma glucose

Body weight growth curves, body composition, and plasma glucose levels were assessed in Gcg-Cre rats of both sexes derived from litters maintained in the FSU colony on the SD lineage (i.e., without crossing to the HsdSage:LE-Rosa26^*tm1(tdTomato)Sage*^ reporter line described in *2.4*).

#### 2.3.1 Body weight growth curves

Body weights were recorded in group- or pair-housed male and female WT, Het, and Homo Gcg-Cre rats in the FSU colony from 4-16 weeks of age (N ≥ 3 per sex/genotype at each age). Body weight curves also were tracked weekly in genotyped Gcg-Cre rats born at the Janvier facility, beginning at 6 wks of age [male: n=17 WT, 30 Het, and 18 Homo; female: N=13 WT, 28 Het, and 22 Homo).

#### 2.3.2 Body composition

Whole body composition (i.e., fat and lean mass) was assessed at a single timepoint in age-matched adult rats from the FSU colony (19-20 wks old; N=4-7 per sex per genotype) using an EchoMRI™ Body Composition Analyzer (Houston, TX).

#### 2.3.3 Plasma glucose levels

In additional adult WT, Het, and Homo Gcg-Cre rats from the FSU colony (15-40 wks old), plasma glucose was assessed under fed and fasting conditions using small blood samples and an EVENCARE G3 Blood Glucose Monitoring System. For this, rats were removed from their home cage between 0900-1000h and gently restrained while a razor blade was used to nick the tail tip. Tail blood (10-15 μl) was collected directly onto a glucose test strip within 15s of each rat’s removal from its home cage. Samples were obtained from rats under conditions of *ad libitum* food access, and from rats deprived of food (but not water) for 21h (including overnight) prior to blood collection.

### 2.4 Generation of Gcg-Cre/tdTom reporter rats

Female Homo Gcg-Cre rats in the FSU colony were bred with male Homo HsdSage:LE-Rosa26^*tm1(tdTomato)Sage*^ rats [Envigo; Long Evans (LE) genetic background] to generate Het Gcg-Cre/tdTom reporter progeny. Some neonatal Gcg-Cre/tdTom pups were anesthetized and perfused on postnatal day (P)2 or P10 (see *2.9*, below) to reveal tdTom reporter labeling during early development, while others were perfused as adults and used for immunolabeling, Cre-dependent viral tracing, and RNAscope fluorescent *in situ* hybridization (FISH; see *2.9 -2.11*, below). Due to their mixed SD/LE genetic background, Gcg-Cre/tdTom reporter rats were not used for chemogenetic behavioral experiments.

### 2.5 Analysis of mRNA expression and protein levels in Gcg-Cre rats

#### 2.5.1 qRT-PCR

To measure *Gcg* and *iCre* mRNA expression within the cNTS, fresh frozen brains from adult WT (N=5 males), Het (N=4 males, 1 female) and Homo (N=5 males) Gcg-Cre rats from the Janvier colony were cut into 50 μm coronal slices using a cryostat (Leica CM3050S). The cNTS was then microdissected using biopsy punches (Integra Miltex, 1 mm diameter) in hindbrain sections corresponding to Bregma -14.5 to -13.3 mm. RNA from brain micropunches was extracted using Trizol (Invitrogen) according to the manufacturer’s protocol, and quantified with Nanodrop (Thermofisher). Reverse transcription was performed using iScript cDNA kit (Biorad) with 1.5 μg of starting RNA. qPCR was performed on a QuantStudio 7 Flex Real-Time PCR System (Applied Biosystems) using TaqMan assays (Thermofisher: *Gapdh* Rn01775763_g1; *Gcg* Rn00562293_m1 and *iCre* Custom assay w2001517107000). Results were quantified using the 2-ddCtt method.

#### 2.5.2 GLP1 protein levels in brain and plasma

WT, Het, and Homo Gcg-Cre rats from the FSU colony were rapidly anesthetized by isoflurane inhalation (5% in oxygen) and then decapitated. Brains were extracted within 1min, separated into brainstem and forebrain portions, snap-frozen in isopentane on dry ice, and then stored at -80°C until protein extraction. Brain tissues were homogenized in ice-cold homogenization buffer for 2-3min using a probe sonicator (Sonic Dismembrator Model 100, Fisher Scientific). Homogenization buffer contained 0.05M phosphate-buffered saline (PBS) with 0.2% (v/v) Triton X, 1.0% (v/v) mammalian protease inhibitor cocktail (Sigma-Aldrich P8340) and 1.0% (v/v) phosphatase inhibitor cocktail #2 and #3 (Sigma-Aldrich P5726, P0044). Tissue homogenates were centrifuged at 14,000×g at 4°C for 20min and supernatants collected and stored at -80°C before initiating the protein assay.

To assay plasma GLP1 levels, blood was collected by either cardiac puncture using K3EDTA blood collection tubes (Fisher Scientific, 22-040-035) immediately before perfusion, or by tail bleeding using SAFE-T-FILL capillary collection tubes (RAM Scientific, 07 6013). Blood samples were collected on ice and centrifuged within 1h at 4°C for 10min at 3000 RPM (Eppendorf, Centrifuge 5804R). Separated plasma was stored at -80°C until the protein assay was initiated.

Total GLP1 protein in brain tissue or in plasma was determined using GLP-1 Total ELISA 96-Well Plate Assay (Sigma-Aldrich, EZGLP1T-36K). The assay was conducted according to manufacturer’s protocol, except that the matrix buffer supplied in the ELISA kit was replaced by homogenization buffer for brain tissue samples. Total protein in brain tissue was determined using a protein assay kit (Pierce™ BCA Protein Assay Kit; ThermoFisher, 23225) following the manufacturer’s protocol.

### 2.6 Microinjection of Cre-dependent AAV to label the axonal projections of Cre-expressing neurons

#### 2.6.1 cNTS-targeted injections

Adult male and female Gcg-Cre rats from the FSU colony (N=4) were anesthetized by isoflurane inhalation (1–3% in oxygen; Halocarbon Laboratories) and placed into a stereotaxic device. The head was ventroflexed by 20 degrees using an adjustable gas anesthesia adaptor (Model 929-B Rat Gas Anesthesia Head Holder). The skin on top of the neck was shaved, disinfected, and incised down the midline (0.75-1.0 cm) just caudal to the occipital bone. The neck muscles were then incised down the midline, bluntly dissected laterally, and held in place with surgical hooks to reveal the dura overlying the cisterna magna. The dural layer was scraped clean using a dental plaque remover, incised using a sterile needle, and reflected to visualize obex. Cre-dependent AAV [i.e., either AAV1-EF1a-double floxed-hChR2(H134R)-EYFP-WPRE-HGHpA, Addgene 20298 or AAV1-pCAG-FLEX-EGFP-WPRE, Addgene 51502] was delivered bilaterally into the cNTS (0.3mm lateral, 0.5mm ventral to obex) by pressure (500 nl/injection, 500 nl/min) using a pulled glass micropipette tip (∼20 μm outer tip diameter) connected to a 10 μl Hamilton syringe, with injection speed and volume controlled by a digital stereotaxic microinjector (Quintessential Stereotaxic Injector, QSI, Cat# 53313). The micropipette was left in place for 3min after each injection. After the injector was removed, the dorsal neck muscles were sutured, and then the overlying skin was sutured. Rats were injected subcutaneously with ketofen (2 mg/kg BW) and buprenorphine (0.03 mg/kg BW) and then returned to their home cages after full recovery from anesthesia. Rats were perfused 3-5 weeks later (see *2.9*, below).

#### 2.6.2 Amygdala-targeted injections

As described in the Results (see *3.4*, below), robust tdTom reporter labeling was observed within the basolateral amygdala (BLA) and other forebrain regions in neonatal and adult Gcg-Cre/tdTom rats. To test whether the unexpected pattern of reporter expression reflected developmentally transient or persistent neural expression of *Gcg* and *iCre*, Cre-dependent AAV1-EF1a-double floxed-hChR2(H134R)-EYFP-WPRE-HGHpA (Addgene 20298) was delivered bilaterally into the BLA in two adult female Gcg-Cre/tdTom rats from the FSU colony. For this, rats were anesthetized by isoflurane inhalation and placed into a stereotaxic device. The shaved skin on top of the skull was disinfected and incised down the midline (0.75-1.0 cm). The skin was reflected to visualize bregma and lambda, and the head adjusted to achieve a flat-skull position. A small hole was drilled through the skull over each BLA injection site. A pulled glass micropipette containing AAV was lowered to the targeted coordinates (from bregma: LM: +/- 4.9 mm, AP: -2.0 mm, DV: -8.0 mm) and delivered as described above *(2.6.1)*. Rats were perfused 3-5 weeks later (see *2.9*, below).

### 2.7 Validation of Cre-dependent excitatory DREADD expression in Gcg-Cre rats

Adult female Het Gcg-Cre rats from the FSU colony (N=14) received bilateral cNTS-targeted delivery of a virus expressing Cre-dependent excitatory DREADD [AAV8-hSyn-DIO-hM3D(Gq)-mCherry; Addgene 44361], using the surgical and viral delivery approach described above *(2.6.1)*. Beginning 3 weeks later, rats were used in a pilot behavioral experiment that is not part of the present report. At least 1 week after completing that experiment, rats were injected i.p. between 0900-0945h with 0.15M NaCl saline vehicle (1.0ml/kg BW) or vehicle containing clozapine-n-oxide (CNO; 1.0mg/ml/kg BW), returned to their home cages, then anesthetized and perfused with fixative 90min later (see *2.9*, below) for subsequent immunocytochemical detection of CNO-induced nuclear cFos and cytoplasmic viral reporter labeling.

### 2.8 Analysis of food intake in Gcg-Cre rats after chemogenetic stimulation of cNTS neurons

#### 2.8.1

In adult male and female Gcg-Cre Het rats from the Janvier colony that were shipped to the Univ. of Gothenburg (Hets: N=8 male, 7 female; WT controls: N=10 male, 7 female), AAV2-hSyn-DIO-hM3D(Gq)-mCherry; Addgene 44361) was delivered bilaterally into the cNTS (0.5μl; 0.1μl/min) through a cannula implanted with tip positioned at the following coordinates: at the level of the occipital suture, ±0.7mm from midline; 5.9mm ventral from skull. After AAV delivery, in the same surgical session, rats were implanted unilaterally with a lateral ventricle (LV) stainless steel guide cannula with the tip positioned at the following coordinates: -0.9mm from bregma; ±1.6mm from midline; 2.5mm ventral from skull. LV cannula placement was confirmed one week after the surgery by measuring water intake in rats after LV infusion of angiotensin II (20ng/2μl) [30]. Only rats that drank at least 5 ml of water in 30min were included in the pharmacological GLP-1R blockade experiment described below (see *2.8.1.2*), which required drug or vehicle delivery through the LV cannula. Correct AAV targeting of the cNTS was confirmed at the termination of the behavioral experiments described below (see *2.8.1.1, 2.8.1.2*). For this confirmation, anesthetized rats were decapitated, brains were collected and flash frozen, the caudal medulla was sectioned on a cryostat, and cannula tracks and Cre-mediated AAV reporter expression was confirmed using a fluorescence microscope, with cNTS reporter labeling confirmed in Het but not in WT rats.

##### 2.8.1.1 Food intake and food choice in male and female rats after cNTS-administered CNO

Three weeks after cannula placement and AAV delivery into the cNTS (described above, *2.8.1*), Gcg-Cre WT and Het rats were pre-exposed to peanut butter to reduce neophobia during a subsequent food choice test. Rats that did not consume any peanut butter during the first exposure received a second exposure; all rats consumed the peanut butter on either their first or second exposure. Prior to the choice test, food was withheld during the second half of the light cycle. Rats were unilaterally injected through the cNTS cannula with CNO [0.5μg in 0.3μl of 1% DMSO in artificial cerebrospinal fluid (ACSF) vehicle] or vehicle alone just before dark onset. Rats were then offered pre-weighed amounts of chow and peanut butter simultaneously in their home cage, with cumulative intake measured manually 1h and 16h post injection. Rat body weights were assessed before and after the 16h food access period. The choice test was repeated 48h later using a within-subjects crossover design, such that each rat was injected into the cNTS with vehicle or CNO in a counterbalanced fashion prior to the food choice test.

##### 2.8.1.2 CNO-induced hypophagia after central GLP1R blockade in male rats

A significant effect of central chemogenetic stimulation of cNTS neurons to reduce chow intake was found in male Het Gcg-Cre rats in the food choice experiment described above *(2.8.1.1)*, with no effect on food choice or peanut butter intake in Het males, and no significant effects on food intake in Het females or in WT rats (see *Results, 3.7.1*). Thus, only Het male rats with Ang II-confirmed LV cannula placements (N=8) were used to determine whether central (LV) delivery of a specific GLP1R antagonist (Exendin 9, EX9) would block the effects of LV-delivered CNO on chow intake and body weight. For this experiment, rats received only 50% of their normal overnight chow consumption on the night prior to each feeding test, in order to increase intake during the test. Before dark onset, each food-restricted rat was injected with EX9 (20μg/1μL in ACSF) or vehicle through the LV guide cannula. Ten minutes later, CNO (1μg/1μL in 1% DMSO in ACSF) or vehicle was injected through the LV guide cannula. Chow was then returned, and cumulative intake measured after 1 and 24h. Body weights were assessed at the onset and termination of each 24h chow intake period. Using a within-subjects design, each rat received each treatment combination across 4 separate chow intake sessions that were spaced 48h apart.

#### 2.8.2 Chemogenetic stimulation using systemic CNO

Food intake was assessed in adult male Het Gcg-Cre rats from the FSU colony (N=14) that received bilateral cNTS-targeted delivery of excitatory DREADD-expressing virus [AAV8-hSyn-DIO-hM3D(Gq)-mCherry; Addgene 44361] at least 3 weeks earlier. Rats were individually housed in standard-sized tub cages with wood chip bedding and wire cage tops, with cages connected to a computerized BioDAQ feeding monitoring system (Research Diets). Rats had *ad libitum* access to pelleted chow (Purina 5001) and water, except as noted below. The food hopper was positioned just outside a small head port opening on one end of the cage to provide oral access to chow through slots in the hopper while restricting the rat’s ability to remove large pieces. The hopper was connected to a digital scale linked to the BioDAQ system to record the time and amount of food removed. Unconsumed chow crumbs fell into a small pan below the hopper (out of the animal’s reach), with spillage weight detected on the same digital scale. Rats were acclimated to the BioDAQ cage system for 7-14d. On at least 5 of these days, port access to food was blocked for 60min before lights out, and rats were injected with 0.15M NaCl (1.0 ml, i.p.) during this 60min period. Food access was returned just before lights out.

After acclimating rats to BioDAQ housing conditions and i.p. injections, a counterbalanced within-subjects experiment was conducted to assess dark-onset food intake in rats receiving systemic CNO (Tocris; 1.0 mg/kg BW, i.p.) alone, or CNO followed by a low (i.e., behaviorally subthreshold) systemic dose of cholecystokinin octapeptide (CCK; Bachem; 1.0μg/kg BW, i.p.). Using a Latin square design, each rat received four treatment combinations in randomized order across four treatment days that were spaced 48-72h apart, as follows: 1) vehicle (0.15M NaCl) followed 30min later by vehicle; 2) vehicle followed 30min later by CCK; 3) CNO followed 30min later by vehicle; and 4) CNO followed 30min later by CCK. Each injection was delivered i.p. in a volume of 1.0ml/kg BW. On treatment days, head port access to food was blocked 2h before dark onset. The first i.p. injection was delivered 35min before dark onset. The second i.p. injection was given 30min later, with food access restored immediately (i.e., within a few minutes of dark onset). Food intake was then recorded over the 12h dark period, with data subsequently binned into 1, 2, 4, 6, and 12h timepoints for statistical analysis. After concluding the experiment, rats were perfused with fixative (*2.9*, below) for immunocytochemical detection of caudal brainstem neurons expressing mCherry viral reporter.

### 2.9 Perfusion fixation, tissue sectioning, and CLARITY

Rats from the FSU colony were anesthetized with a lethal dose of pentobarbital sodium (Fatal Plus or Euthasol, 130mg/kg; Butler Schein, Columbus, OH or Covetrus, Dublin, OH) and then transcardially perfused with a physiological saline rinse followed by 4% paraformaldehyde in 0.1M sodium phosphate buffer. Brains were extracted from the skull, postfixed overnight, and then cryoprotected in 20% sucrose solution for 24–72 h at 4°C. Adult rat brains were sectioned coronally, horizontally, or sagitally (35μm) using a freezing sliding microtome; brains from P2 and P10 pups were sectioned coronally (50μm). Sections were collected sequentially into 4-6 adjacent sets and stored in cryopreservant solution [31] at −20°C for subsequent processing.

To visualize tdTom-positive cells in pancreatic and intestinal tissue, neonatal (P2, P10) Gcg-tdTom pups (N=4) were anesthetized with pentobarbital sodium and transcardially perfused with saline followed by ice-cold hydrogel solution (4.0% PF, 4.0% acrylamide, 0.02% bis-acrylamide and 0.25% VA-044 initiator in 0.1M PBS), according to our published protocol [32]. The pancreas and small intestine were dissected out of the abdominal cavity. The pancreas was postfixed in hydrogel solution for 24-36h at 4°C. The small intestine was opened longitudinally along the mesentery side, rinsed briefly with ice-cold saline to remove gut contents, and then postfixed in hydrogel solution for 24-36h at 4°C. Postfixed tissues were rinsed briefly with saline, patted dry with a Kimwipe to remove excess hydrogel solution, submerged in 100% mineral oil and polymerized for 3h at 37°C. Removing excess hydrogel solution on the tissue surface and replacing hydrogel solution with mineral oil as the polymerization medium served two purposes: 1) to separate hydrogel-infused tissue from oxygen in the air during polymerization; and 2) to eliminate the need to separate polymerized tissue from polymerized hydrogel within the polymerization medium, thereby reducing damage to the tissue samples. Polymerized tissues were rinsed 3-4 times with 0.5M borate buffer (BB) to remove excess mineral oil, and then passively cleared in clearing solution (4% SDS in 0.5M BB, pH 8.5) until clear, which took 3-10 days depending on the tissue sample. Cleared tissue was rinsed in 0.5M BB for 24h with 4-5 changes to remove excess SDS, and saved in BB at room temperature until immunofluorescent labeling procedures were initiated.

### 2.10 RNAscope-based FISH

Manual assay probes and detection reagents were purchased from Advanced Cell Diagnostics (ACD). *Gcg* and *iCre* mRNA were identified using probes for Rn-Gcg (315471-C2, Accession No. NM_012707.2) and iCre (423321-C2, Accession No. AY056050.1), and by using the RNAscope® Multiplex Fluorescent Reagent Kit v2 (323110). Selected tissue sections were removed from cryoprotectant and washed for 1h in six changes of 0.1M Phosphate Buffer (PB). Sections were pretreated at room temperature with H_2_O_2_ solution (ACD 322335) for 30min, followed by six 5min rinses in 0.1M PB. Sections were mounted out of 0.01M Tris Buffer (TB) onto Epredia™ Gold Seal™ UltraStick™ Adhesion Microscope Slides (Thermo Fisher Scientific, USA). Slides were baked at 60°C in the HybEZTM oven for 1h, dipped into 100% ethanol for 10s and then air-dried for 15min before a hydrophobic barrier was drawn around tissue sections using an ImmEdge® Pap Pen (Vector Laboratories Inc. H-400). After air-drying overnight, slides were either stored in a plastic box in a -20°C freezer for later processing, or were used immediately. Incubations were performed at 40°C within a HybEZTM Oven, using the HybEZTM Humidity Control Tray unless otherwise noted. Two to four drops of each reagent solution were used to cover tissue sections, followed by three 3min rinses in 1X RNAscope® Washing Buffer (ACD 320058) at room temperature. Sections were then incubated with Protease IV (ACD 322336) for 20-25min at room temperature, followed by three dips in distilled H_2_O and two 2min rinses in 0.01M TB. Sections were then incubated in Rn-Gcg-C2 (1:50) or iCre-C2 probes (1:50 in probe diluent) followed sequentially by amplification steps with Amp1 (ACD 323101) for 30min, Amp2 (ACD 323102) for 30min, and Apm3 (ACD 323103) for 15min. *Gcg* or *iCre* mRNA transcripts were labeled with fluorophore-conjugated Tyramine plus (TSAP). Sections were incubated in channel 2-specific HRP (ACD 323105) for 15min followed by Cy5-TSAP (1:2K for Gcg, 1:1K for iCre) for 30min. After the final washing buffer rinse, sections were incubated in DAPI (ACD 323108) at room temperature for 1min before coverslipping using Fluoromount-G (SouthernBiotech, 0100-01).

### 2.11 Immunolabeling

#### 2.11.1 Brain Sections

Tissue sections were removed from cryopreservant storage, rinsed in 0.1M sodium phosphate buffer, and pretreated for 20min in 0.5% sodium borohydride and 15min in 0.15% H_2_O_2_ with an intervening 30min rinse in 0.1M PB. Primary and secondary antisera were diluted in buffer containing 0.3% Triton X-100 and 1% normal donkey serum. Tissue sections were rinsed for 30min in several changes of buffer between incubation steps. All primary and secondary antibodies, reagents, and working dilutions are listed in **Table 2**. For dual immunoperoxidase localization of cFos and Cre-dependent mCherry viral reporter, tissue sections were incubated in a rabbit monoclonal antiserum against cFos followed by biotinylated donkey anti-rabbit IgG. Sections were then treated with Elite Vectastain ABC reagents (Vector) and reacted with nickel sulfate, diaminobenzidine (DAB), and H_2_O_2_ to produce a blue-black nuclear cFos reaction product. Next, to enhance visualization of mCherry, cFos-labeled tissue sections were incubated in guinea pig polyclonal antiserum against RFP, followed by incubation in biotinylated donkey anti-guinea pig IgG and Elite Vectastain ABC reagents, with a final reaction using plain DAB and H_2_O_2_ to produce a brown cytoplasmic reaction product. Immunolabeled tissue sections were mounted onto SuperFrost plus microscope slides, dehydrated in ascending ethanol solutions, defatted with xylene, and coverslipped using Cytoseal 60 (VWR).

**Table 2.** Viruses, antibodies, reagents, and incubation parameters.

| Name | Source | Incubation | Experiment |
| --- | --- | --- | --- |
| AAV-hSyn-DIO-hM3D(Gq)-mCherry | Addgene, #44361-AAV8 |  | Chemogenetic stimulation for feeding behavior, cFos |
| AAV8-hSyn-DIO-mCherry | Addgene #50459-AAV8 |  | Control virus for chemogenetic stimulation, cFos |
| Alexa Fluor® 488 Donkey Anti-Rabbit IgG | Jackson ImmunoResearch, 711-545-152 | 1:500, 3h, RT | Immunofluorescent labeling |
| Alexa Fluor® 647 Donkey Anti-Rabbit IgG | Jackson ImmunoResearch, 711-605-152 | 1:500, 3h, RT | Immunofluorescent labeling |
| Biotin-SP Donkey Anti-Rabbit IgG | Jackson ImmunoResearch, 711-065-152 | 1:1K, 2h, RT | Immunoperoxidase labeling |
| Chicken Anti-Green Fluorescent Protein | Abcam, ab13970, RRID:AB_300798 | 1:1K, 24h, RT | Immunofluorescent labeling |
| Cy™3 Donkey Anti-Guinea Pig IgG (H+L) | Jackson ImmunoResearch, 706-165-148 | 1:300, 40h, 37°C | CLARITY |
|  |  | 1:500, 3h, RT | Immunofluorescent labeling |
| Donkey anti-Rabbit IgG, HRP conjugated | Millipore Sigma, AP 182P | 1:500, 40h, 37°C | CLARITY |
|  |  | 1:500, 3h, RT | TSAP-enhanced Immunofluorescent labeling |
| Guinea Pig Anti-Red Fluorescent Protein | Synaptic Systems, 390 004, RRID:AB_2737052 | 1:15K, 48h, RT | Immunofluorescent labeling |
|  |  | 1:10K, 24h, RT | Immunoperoxidase labeling |
| *iCre* mRNA | Advanced Cell Diagnostics, 423321-C2, Accession No. AY056050.1 | 1:50, 2h 40°C | RNAscope labeling |
| Mineral oil | VWR Life Science, J217 | 100%, 3h, 37°C | CLARITY |
| Mouse Anti-Dopamine $\beta$ Hydroxylase | Millipore Sigma, MAB308, RRID:AB_2245740 | 1:8K, 48h, RT | Immunofluorescent labeling |
| Rabbit Anti c-Fos (9F6) | Cell Signaling, #2250, RRID:AB_2247211 | 1:10K, 24h, RT | Immunoperoxidase labeling |
| Rabbit Anti-Glucagon-Like Peptide 1 (7-37) | Peninsula Laboratories, T-4363, RRID:AB_518978 | 1:10K, 90h, 37°C | CLARITY |
|  |  | 1:25K, 48h, RT | TSAP-enhanced Immunofluorescent labeling |
| Rn-*Gcg* mRNA | Advanced Cell Diagnostics, 315471-C2, Accession No. NM_012707.2 | 1:50, 2h 40°C | RNAscope labeling |
| TSA™Plus Cyanine 5 | AKOYA Biosciences,<br>SAT705A001KT | Clarity: 1:400, 40h, 37°C | CLARITY |
|  |  | 1:1K & 1:2K, 30min, 40°C | RNAscope labeling |
| 2,20-Thiodiethanol | Sigma-Aldrich, 166782 | 15%, 30%, 45%, 60%, 5h<br>each, RT | CLARITY |
| 2% bis-Acrylamide | RPI Research Products,<br>A11275 |  | CLARITY |
| 40% Acrylamide<br>Solution | RPI Research Products,<br>A11265 |  | CLARITY |

#### 2.11.2 CLARITY-processed pancreatic and intestinal tissues

Details regarding antibodies, reagents, and labeling conditions are summarized in **Table 2**. Primary and secondary antibodies were diluted using 0.5M BB (pH 8.5) containing 1% donkey serum and 0.3% triton X. Cleared tissue samples were either processed for Refractive Index Matching (RIM) in ascending TDE solutions to visualize Gcg in native tdTom form, or were first processed for GLP1 immunofluorescent labeling to achieve simultaneous visualization of GLP1 and native tdTOM. For GLP1 labeling, the pancreas or a segment of small intestine was incubated sequentially in rabbit anti-GLP1, HRP-conjugated donkey anti-rabbit IgG, and TSAP-conjugated Cy5, as specified in **Table 2**. Each incubation was followed by a 4h wash in 3 changes of BB at 37°C. Labeled tissues were processed for RIM in ascending TDE solutions (**Table 2**), and then stored at room temperature in 60% TDE until imaging.

### 2.12 Microscopic Imaging

#### 2.12.1 Fluorescence and immunoperoxidase imaging

Low magnification images of immunofluorescence or immunoperoxidase labeling were collected using a Keyence (BZ-X700) microscope and 2X objective, with or without tiling. Higher magnification images of immunoperoxidase labeling were collected using a 10X or 20X objective. Collected single and tiled images were processed using “Full Focus” and “XY-Stitching” features of the Keyence Analyzer BZ-H4XD software to obtain Z-max projection images.

#### 2.12.2 Image acquisition and quantification of FISH (RNAscope) labeling

High-resolution images were acquired using a Leica TCS SP8 Confocal Microscope with a 100× oil objective. Cy5 and DAPI were excited using a 638 nm and 405 nm Diode laser, respectively. One to two rostro-caudally matched medullary sections were imaged in each rat. For each image, twenty 0.28μm-thick optical sections were collected through the cNTS and a Z-max projection image was generated using Leica LAS version 4.0 image software. Labeled *Gcg* mRNA transcripts were quantified from these Z-max projection images using the HALO multiplex FISH module (FISH-IF V2.1.7, Indica Labs, Albuquerque, New Mexico USA), designed and optimized for FISH RNAscope quantification. Briefly, cell boundaries were identified by nuclear segmentation based on DAPI staining and parameters of maximum cytoplasm detection. *Gcg* mRNA-positive neurons were detected using parameters for Cy5 threshold (nucleus, cytoplasm). Each labeled *Gcg* mRNA transcript was defined by spot segmentation (threshold, size, intensity, segmentation aggressiveness). When *Gcg* labeling was too intense to be separated into individual transcripts (primarily in WT rats), Gcg mRNA transcript copy number within the cell was automatically calculated based on labeling intensity and area. All parameters were adopted from the HALO multiplex FISH module with subtle fine-tuning based on *Gcg* mRNA labeling across tissue samples; importantly, images were analyzed using identical settings across all genotypes. In each animal, the average mRNA copy number per *Gcg*-positive cell was exported from HALO and used to compare the quantity of *Gcg* mRNA transcripts within the cNTS across genotypes.

#### 2.12.3 Imaging of CLARITY-processed pancreatic and intestinal tissues

Images were acquired using a Leica TCS SP8 confocal microscope equipped with a 25X CLARITY objective (6.0-mm working distance; HC FLUOTAR L 259/1.00 IMM; ne = 1.457; motCORR VISIR; Leica) designed to match the 1.45 refractive index of cleared hybridized tissue. Labeled tissues were positioned in a 60×15mm clear polystyrene tissue culture dish (Fisher Scientific, 08-772B) filled with fresh 60% TDE solution. The CLARITY objective was then submerged directly into the TDE solution, as previously described [32]. Native tdTom was excited using a 552 nm OPSL laser; Cy5-labeled GLP1 was excited using a 638 nm Diode laser. Optical sections were collected and Z-max and/or 3D projections were generated using Leica LAS 4.0 image software.

### 2.13 Data Analysis

Group summary data in text and graphs are expressed as mean ± SEM, with individual data points shown. Statistical analyses were conducted using Prism 9. BW, fat mass, and lean mass were compared using 2-way ANOVA, with genotype and sex as between-subjects factors. Plasma glucose levels were compared using multivariate (3-way) ANOVA, with genotype, sex, and feeding status as between-subjects factors. Hindbrain *Gcg* mRNA labeling was analyzed using 1-way ANOVA, with genotype as the between-subjects variable. Total GLP1 protein levels in brain tissue were analyzed using 2-way ANOVA, with genotype as the between-subjects variable and brain region as the within-subjects variable. Plasma GLP1 levels were analyzed using 1-way ANOVA, with genotype as the between-subjects variable. Intake data from the chow vs. peanut butter choice experiment were analyzed using 2-way ANOVA with sex and CNO as factors. Chow intake data from the Ex9 blockade experiment were analyzed using 2-way ANOVA with CNO and Ex9 treatments as between-subject factors. Chow intake in the i.p. CNO/CCK experiment was analyzed using multivariate ANOVA, with four i.p. injection conditions and time as within-subject factors. Counts of DREADD viral reporter- and/or cFos-positive cells were analyzed using 2-way ANOVA, with i.p. injection before perfusion and AAV as between-subject variables. For each analysis, significant main effects or interactions were followed by post-hoc t-tests corrected for repeated comparisons when appropriate.

## 3.0 Results

### 3.1 General health, growth, body composition, and plasma glucose levels of Gcg-Cre rats

Male and female WT, Homo and Het Gcg-Cre rats are fertile when bred with local colony-derived WT Gcg-Cre or outbred SD rats (Envigo). Homo Gcg-Cre female rats also breed successfully with male HsdSage:LE-Rosa26^*tm1(tdTomato)Sage*^ rats, despite different genetic background strains (i.e., SD and LE). Regardless of breeding strategy, Gcg-Cre rat litter sizes average 10.2 ± 0.4 pups (based on more than 50 litters). Pre-weaning pups of each genotype develop normally after birth, displaying typical body sizes and survival rates.

#### 3.1.1 Body weight growth curves

There was no significant effect of genotype on BW assessed from 4-16 weeks of age in male WT, Het, and Homo Gcg-Cre rats in the FSU colony [**Fig. 2A**; F(2,105) = 2.417, *P* = 0.09]. Conversely, there was a significant effect of genotype on BW in female Gcg-Cre rats [**Fig. 2B**; F(2,83) = 14.99, *P* < 0.001]. Post-hoc comparisons confirmed that female Homo Gcg-Cre rats weighed slightly but significantly more than female Het and/or WT rats after 5 weeks of age, whereas the BWs of Het and WT female rats did not differ at any time (**Fig. 2B)**. Similar results were obtained in Gcg-Cre rats that were re-derived and bred in the Janvier facility: genotype had no effect on body weight in males, whereas Homo Gcg-Cre females in the Janvier colony showed a non-significant trend towards weighing more than WT and Het females from 6-9 wks of age [F(2,60) = 2.070, *P* = 0.13; **Fig. S1**].

**Figure 2.**
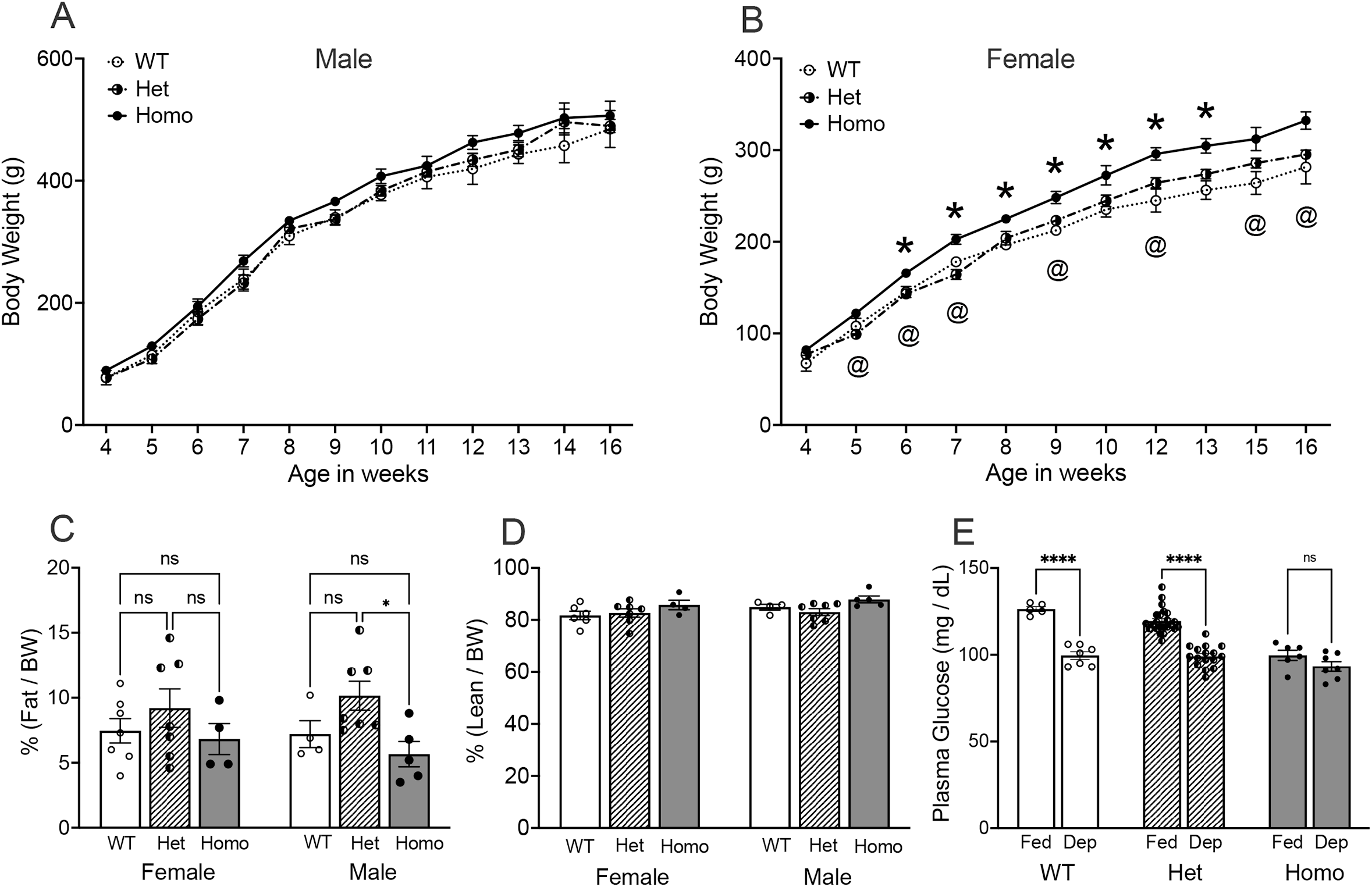
Body weight (BW), body composition, and plasma glucose levels in Gcg-Cre rats from the FSU colony. **A**, Male Gcg-Cre rats display similar growth curves and BWs after weaning, regardless of genotype. **B**, Female Homo Gcg-Cre rats display slightly but significantly higher BW compared to WT rats (\**P* < 0.05, Homo vs. WT) and Het rats (^@^*P* < 0.05, Homo vs. Het). **C**, There is no significant effect of genotype on body fat percentages in female Gcg-Cre rats, whereas Het Gcg-Cre rats have more body fat compared to Homo Gcg-Cre rats (\**P* < 0.05). **D**, Genotype does not impact lean mass in Gcg-Cre rats of either sex. **E**, Morning plasma glucose levels in male and female *ad lib* fed (Fed) and overnight food deprived (Dep) Gcg-Cre rats. Baseline (Fed) plasma glucose levels were lower in Homo Gcg-Cre rats compared to WT and Hets (*P* < 0.05 for each comparison). Plasma glucose levels fell in Dep WT and Het rats compared to baseline (\**P* < 0.001 within each genotype). Conversely, glucose levels in Homo rats did not fall after overnight food deprivation.

#### 3.1.2 Body composition

EchoMRI™ conducted in 19-20 wk old Gcg-Cre rats revealed no main effect of sex [F(1,28) = 0.02, *P* = 0.88] and no significant interaction between sex and genotype on fat mass as proportion of BW [F(2,28) = 0.39, *P* = 0.68]. However, there was a significant main effect of genotype on fat mass [F(2,28) = 4.35, *P* = 0.023]. Post-hoc comparisons revealed no differences in fat mass among female WT, Het, and Homo rats (**Fig. 2C;** all *P* values > 0.39). Among males, however, Het Gcg-Cre rats had higher proportional fat mass compared to Homo rats (*P* = 0.03; **Fig. 2C**), but not compared to WT rats. Regarding lean mass as proportion of BW (**Fig. 2D**), there was no main effect of sex [F(1,28) = 2.09, *P* = 0.16], and no significant interaction between sex and genotype [F(2,28) = 0.47, *P* = 0.63]. While ANOVA revealed a significant main effect of genotype on lean mass [F(2,28) = 3.58, *P* = 0.042], post-hoc comparisons revealed no differences in lean mass among WT, Het, and Homo rats within either sex (**Fig. 2D;** all *P* values > 0.05).

#### 3.1.3 Plasma glucose levels

There was no main effect of sex on plasma glucose levels in adult Gcg-Cre rats [F(1,65) = 0.16, *P* = 0.69], and no interaction between sex and genotype [F(2,65) = 0.79, *P* = 0.46]. Thus, plasma glucose data from both sexes were combined for further analysis, and as depicted in **Fig. 2E**. There were significant main effects of feeding status [F(1,58) = 87.03, *P* < 0.001] and genotype [F(2,58) = 23.49, *P* < 0.001] on plasma glucose levels, and a significant interaction between the two [F(2,58) = 8.15, *P* < 0.001]. Post-hoc tests confirmed that under baseline fed conditions, plasma glucose levels were lower in Homo Gcg-Cre rats compared to WT and Het rats (**Fig. 2E**; *P* < 0.05 for each comparison). Interestingly, while overnight food deprivation predictably reduced plasma glucose in WT and Het Gcg-Cre rats, the hypoglycemic effect of fasting was absent in Homo rats (**Fig. 2E)**.

### 3.2 Efficiency and selectivity of iCre mRNA expression

In Gcg-Cre/tdTom reporter rats, all (100%) tdTom-labeled neurons within the cNTS and IRt were GLP1-immunopositive, and vice-versa (**Fig. 3A-C**). To obtain additional confirmation of the efficiency and selectivity of *iCre* expression, RNAscope-based FISH and immunocytochemical labeling was performed in caudal brainstem tissue sections from Gcg-Cre/tdTom rats to detect *iCre* mRNA transcripts in GLP1-positive, tdTom-labeled neurons within the cNTS and IRt. In both hindbrain regions, all (100%) GLP1/tdTom-positive neurons expressed detectable *iCre* mRNA, and all (100%) *iCre*-expressing cells were immunopositive for both GLP1 and tdTom (**Fig. 3D,E**).

**Figure 3.**
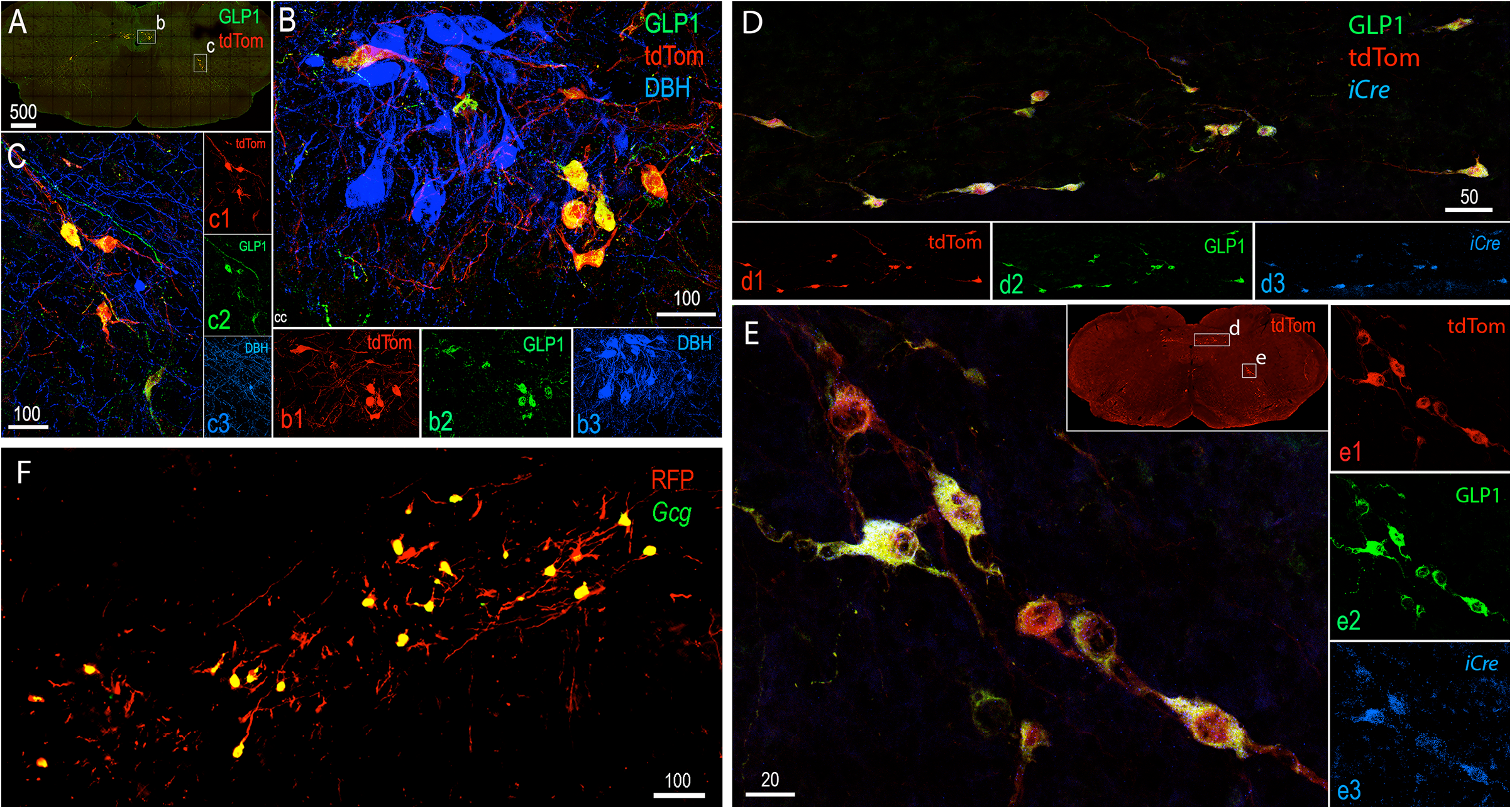
Efficiency and selectivity of hindbrain *iCre* expression in Gcg-Cre/tdTom reporter rats. **A**, low magnification view of the hindbrain distribution of tdTom (red) and GLP1 (green) immunolabeling. The boxed regions (b, c) are shown in higher-magnification confocal images in panels B and C. **B**, Blue (DBH-positive) noradrenergic neurons in the cNTS are not tdTom-positive. All tdTom-positive cNTS neurons are GLP1-positive, and vice-versa. **C**, Blue (DBH-positive) noradrenergic neurons in the IRt are not tdTom-positive. All tdTom-positive IRt neurons are GLP1-positive, and vice-versa. **D**, RNAscope FISH reveals iCre (blue) mRNA expression in tdTom/GLP1 positive neurons within the cNTS (D) and within the IRt (**panel E**). Boxed regions shown in the inset in panel E are shown in higher-magnification confocal images in panels D and E. Scale bars are in microns.

In addition to neurons within the hindbrain cNTS and IRt, a small population of apparently locally-projecting interneurons within the olfactory bulb express *Gcg* in adult rodents, based on *in situ* hybridization in adult rats [6] and fluorescent reporter labeling in adult mice [7][33]. In neonatal and adult Gcg-Cre/tdTom reporter rats, we observed much more extensive olfactory bulb tdTom reporter labeling compared to those published reports, with tdTom-positive cells in Gcg-Cre/tdTom reporter rats including both periglomerular and granule cells (**Fig. 4A,B**). However, based on RNAscope FISH in adult Gcg-Cre/tdTom reporter rats, only a subset of tdTom-positive cells within the olfactory bulb contained detectable *Gcg* mRNA transcripts (**Fig. 4C**). We expand upon this finding in a later section (see *3.4)* as it relates to patterns of additional CNS reporter labeling in neonatal and adult Gcg-Cre/tdTom rats and in a similar transgenic mouse model.

**Figure 4.**
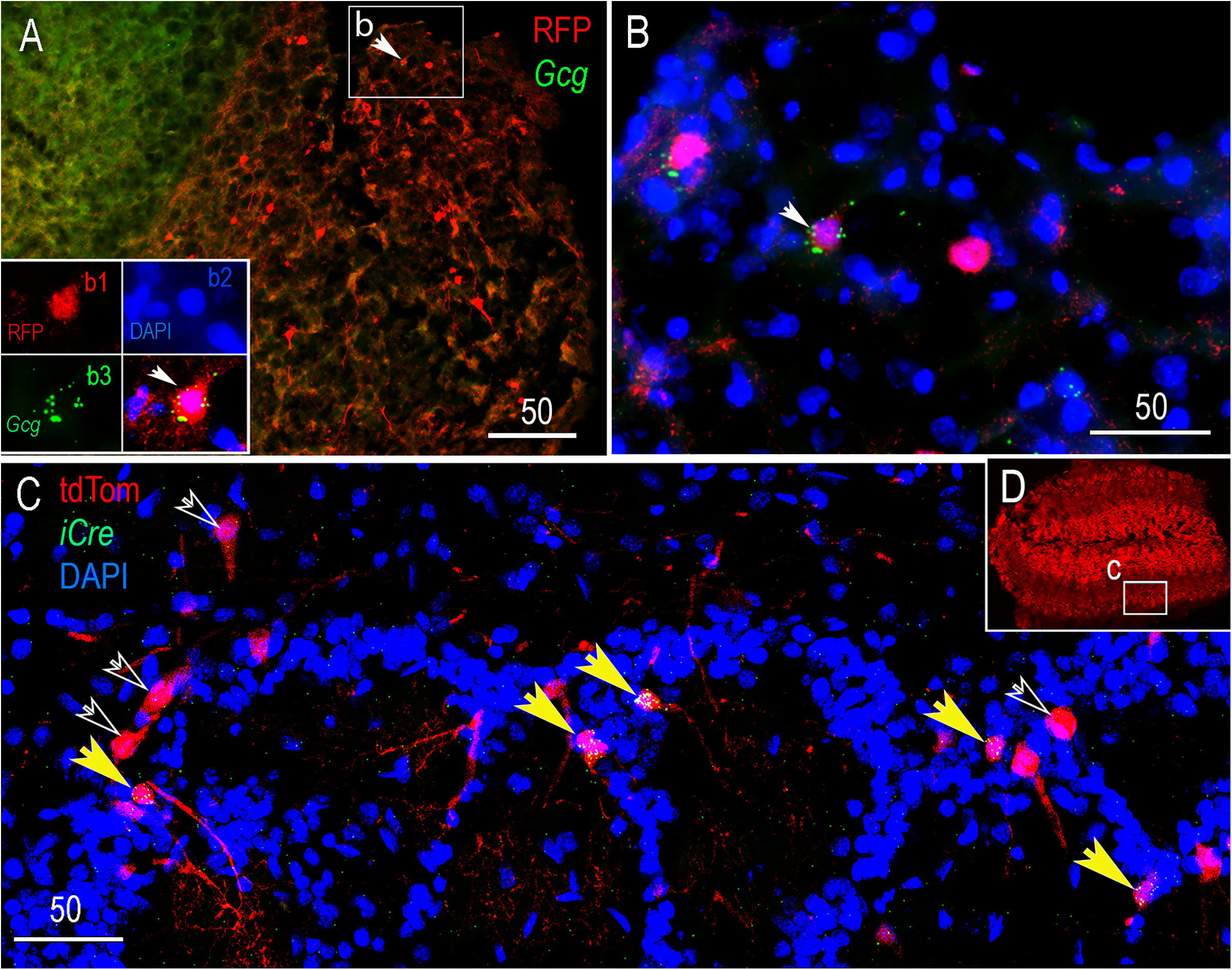
*Gcg* and *iCre* mRNA expression (RNAscope) in the olfactory bulb in adult Gcg-Cre/tdTom reporter rats. **A**, sagittal section through the olfactory bulb. The boxed inset (b) is shown at higher magnification in panel B. Within the inset, a white arrow points out a representative cell immunolabeled for red fluorescent protein (RFP = tdTom-positive). The 4 panels in the lower left of panel A show Gcg mRNA expression (green) in the RFP-positive cell. DAPI nuclear labeling is shown in blue. **B**, *Gcg* mRNA is colocalized to one RFP-positive cell (arrow), but not to two other RFP-positive cells in the same region. **C**, iCre mRNA (green) is colocalized in several tdTom-positive cells within the periglomular olfactory bulb (yellow arrows), but is not detected in other tdTom-positive cells (open arrows). **D**, lower-magnification view of tdTom labeling in the olfactory bulb. The boxed region (c) is shown at higher magnification in panel C. Scale bars are in microns.

### 3.3 Quantitative analysis of Gcg mRNA expression and GLP1 protein levels in Gcg-Cre rats

During microscopic inspection of GLP1 immunocytochemical labeling in Gcg-Cre rats, it became apparent that GLP1 immunolabeling intensity was less robust in Homo compared to WT and Het rats, the latter including Gcg-Cre/tdTom rats (which are Het for the mutant *iCre* knock-in allele). To determine whether cellular expression of *Gcg* mRNA was reduced by the *iCre* knock-in, *Gcg* mRNA was detected using FISH in hindbrain tissue sections from adult male and female Homo, Het, and WT Gcg-Cre rats (N=3-4 per genotype), with transcripts per cell quantified using HALO image analysis software (**Fig. 5**). ANOVA confirmed a significant main effect of genotype on *Gcg* mRNA transcript labeling [F(2,7) = 19.80, *P* = 0.0013]. Post-hoc comparisons indicated that the average number of *Gcg* mRNA transcripts per cell was significantly lower in Homo and Het rats compared to WT rats (*P* < 0.004 for each comparison). Het rats had a higher average level of Gcg mRNA transcript labeling per cell compared to Homo rats (**Fig. 5G**), but this difference was not significant (*P* = 0.41). RNAscope results were extended using qRT-PCR to examine *Gcg* and *iCre* expression in cNTS tissue punches obtained from WT, Het, and Homo Gcg-Cre rats reared in the Janvier colony and shipped to Univ. of Gothenburg. *Gcg* mRNA levels were not different between WT and Het rats, whereas *Gcg* expression in Homo rats was barely detectable (**Fig. 5H**). Het and Homo Gcg-Cre rats displayed similar levels of *iCre* mRNA expression (**Fig. 5H**). *iCre* expression in individual Het rats was highly correlated with *Gcg* expression levels in the same rat (r = 0.99, *P* < 0.0001), but there was no significant correlation in Homo rats (**Fig. 5I**).

**Figure 5.**
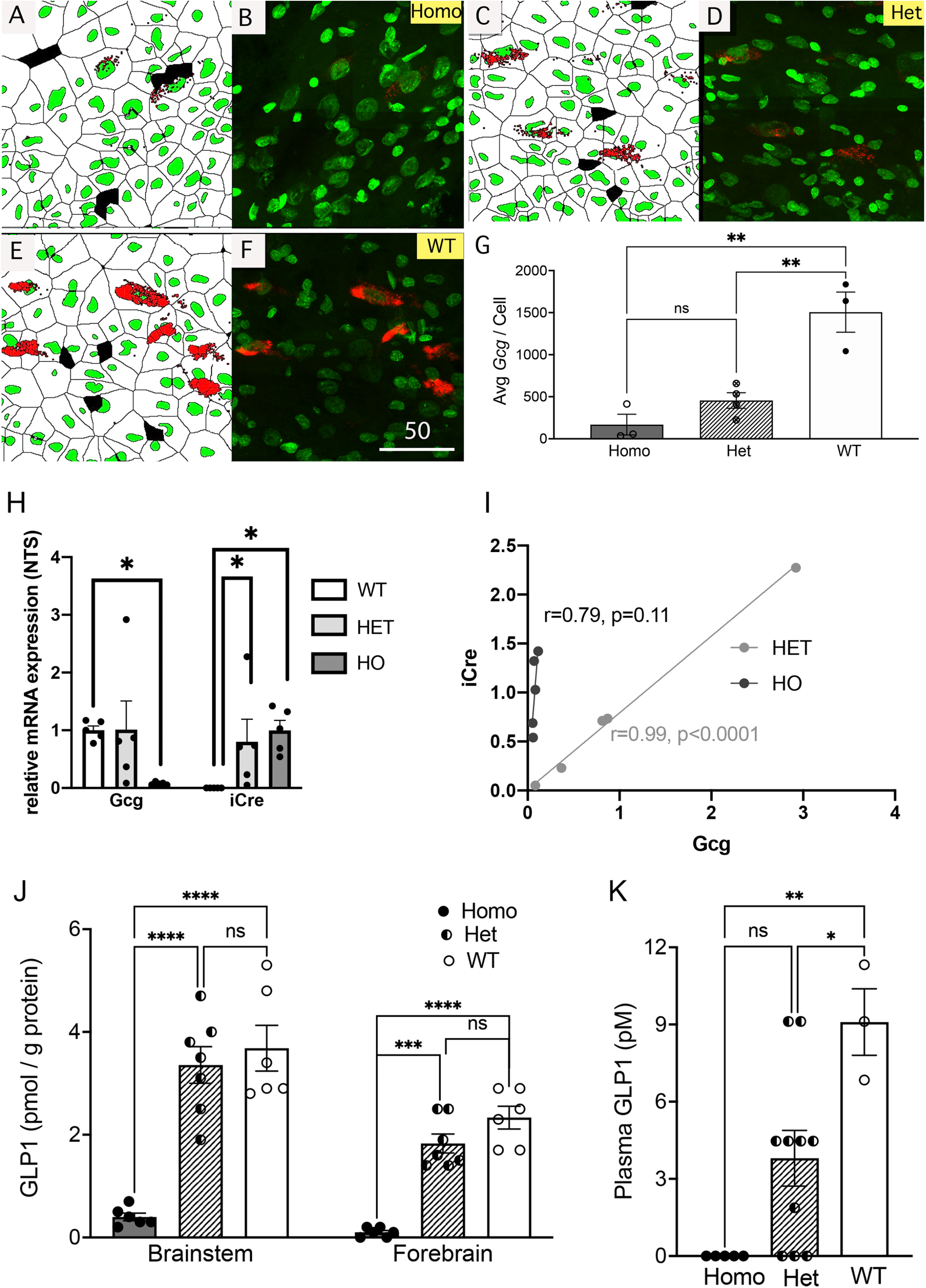
Quantification of *Gcg* mRNA expression and GLP1 protein levels in Gcg-Cre rats. **A-F**, RNAscope FISH followed by HALO-based image analysis of cellular *Gcg* mRNA transcript levels within the cNTS. **A+B**, Homo Gcg-Cre rats display very low cellular levels of Gcg mRNA (red). **C+D**, *Gcg* mRNA transcript labeling (red) in Het Gcg-Cre rats. **E+F**, *Gcg* mRNA transcript labeling (red) in WT Gcg-Cre rats. **G**, summary data depicting the average number of *Gcg* mRNA transcripts quantified in Homo, Het, and WT Gcg-Cre rats. Cellular levels of *Gcg* mRNA in Homo and Het rats are lower than in WT rats (\*\**P* < 0.01 for each comparison). **H**, qRT-PCR analysis of *Gcg* and *iCre* mRNA expression within the NTS. *Gcg* mRNA expression in Het rats (light gray bars) is not significantly different compared to WT rats, whereas *Gcg* mRNA expression in Homo rats (Ho, dark gray) is nearly undetectable. WT rats lack detectable *iCre* expression, whereas *iCre* mRNA levels are similar in Het and Homo rats. **I**, within-subjects correlation of *iCre* and *Gcg* mRNA expression within the NTS in Het and Homo rats. Expression levels are variable but tightly correlated in Het rats, but are not correlated in Homo rats, which have barely detectable levels of *Gcg* mRNA expression. **J**, GLP1 protein levels analyzed via ELISA in tissue samples from the brainstem and forebrain in Gcg-Cre rats. Brainstem and forebrain GLP1 protein levels are similar in Het and WT Gcg-Cre rats, but are markedly lower in Homo Gcg-Cre rats (\*\**P* < 0.001). **K**, Plasma levels of GLP1 protein analyzed via ELISA in Gcg-Cre rats. Compared to plasma GLP1 levels in WT rats, levels in Het rats are reduced by approximately 50%, and levels in Homo rats are essentially undetectable (\**P* < 0.05; \*\**P* < 0.01).

As might be expected, reduced *Gcg* mRNA transcript labeling within the cNTS was associated with markedly reduced levels of GLP1 protein detected by ELISA in homogenized brainstem and forebrain tissue from Homo Gcg-Cre rats (**Fig. 5J;** there was no significant main effect of sex and no interactions; thus, Fig. 5J and K represent combined data from both sexes). ANOVA revealed a significant interaction between brain region (forebrain, brainstem) and genotype on GLP1 protein levels [F(2,16) = 4.46, *P* = 0.03], and significant main effects of brain region [F(1,16) = 34.40, *P* < 0.0001] and genotype [F(2,16) = 45.46, *P* < 0.0001]. Post-hoc comparisons indicated that brainstem and forebrain GLP1 protein levels did not differ between WT and Het Gcg-Cre rats (*P* > 0.3 for each brain region; **Fig. 5J**). Conversely, GLP1 protein levels in the brainstem and forebrain of Homo Gcg-Cre rats were significantly lower compared to levels in both WT and Het rats (*P* < 0.0002 for each genotype comparison for each brain region; **Fig. 5J**).

ANOVA also revealed a significant effect of genotype on plasma GLP1 levels [F(2,15) = 10.12, *P* = 0.002]. Plasma GLP1 was essentially undetectable in Homo Gcg-Cre rats (**Fig. 5K**), in which levels were significantly reduced compared to levels in WT rats (*P* = 0.001). Average plasma GLP1 levels also were lower in Het compared to WT rats (*P* = 0.03), while the difference between Homo and Het rats in plasma GLP1 levels approached but did not reach significance (*P* = 0.06; **Fig. 5K**).

### 3.4 Gcg-Cre/tdTom reporter labeling

#### 3.4.1 Central nervous system (CNS)

The brain distribution of tdTom reporter labeling was identical in neonatal (P1), P10, and adult Gcg-Cre/tdTom rats. Selected regions of CNS labeling in adult Gcg-Cre/tdTom rats are shown in **Figure 6**. Within the hindbrain, tdTom-labeled neurons were located almost exclusively within the cNTS and IRt, with some additional neurons observed in the midline raphe at the same caudal medullary levels (**Fig. 6A,B**). A parasaggital view of tdTom-positive cNTS and IRt neurons reveals their extensive dendritic projections (**Fig. 6C**). At higher levels of the neuraxis, tdTom-labeled cells were present in regions that lack detectable *Gcg* expression in adult rats, including discrete regions of the rostral medulla, cerebellum (**Fig. 6D**), midbrain (**Fig. 6E**), and telencephalon (**Fig. 6F,G**). Scattered tdTom-labeled cells were present within the dorsal cochlear nucleus, cerebellar cortex, inferior colliculus, and perirhinal, insular, and orbital cortices. Larger numbers of tdTom-positive neurons were clustered in the basolateral amygdala (apparent in the inset of **Fig. 6F**), anterior olfactory nucleus, endopiriform nucleus, and the piriform cortex and olfactory bulb (**Fig. 6G**).

**Figure 6.**
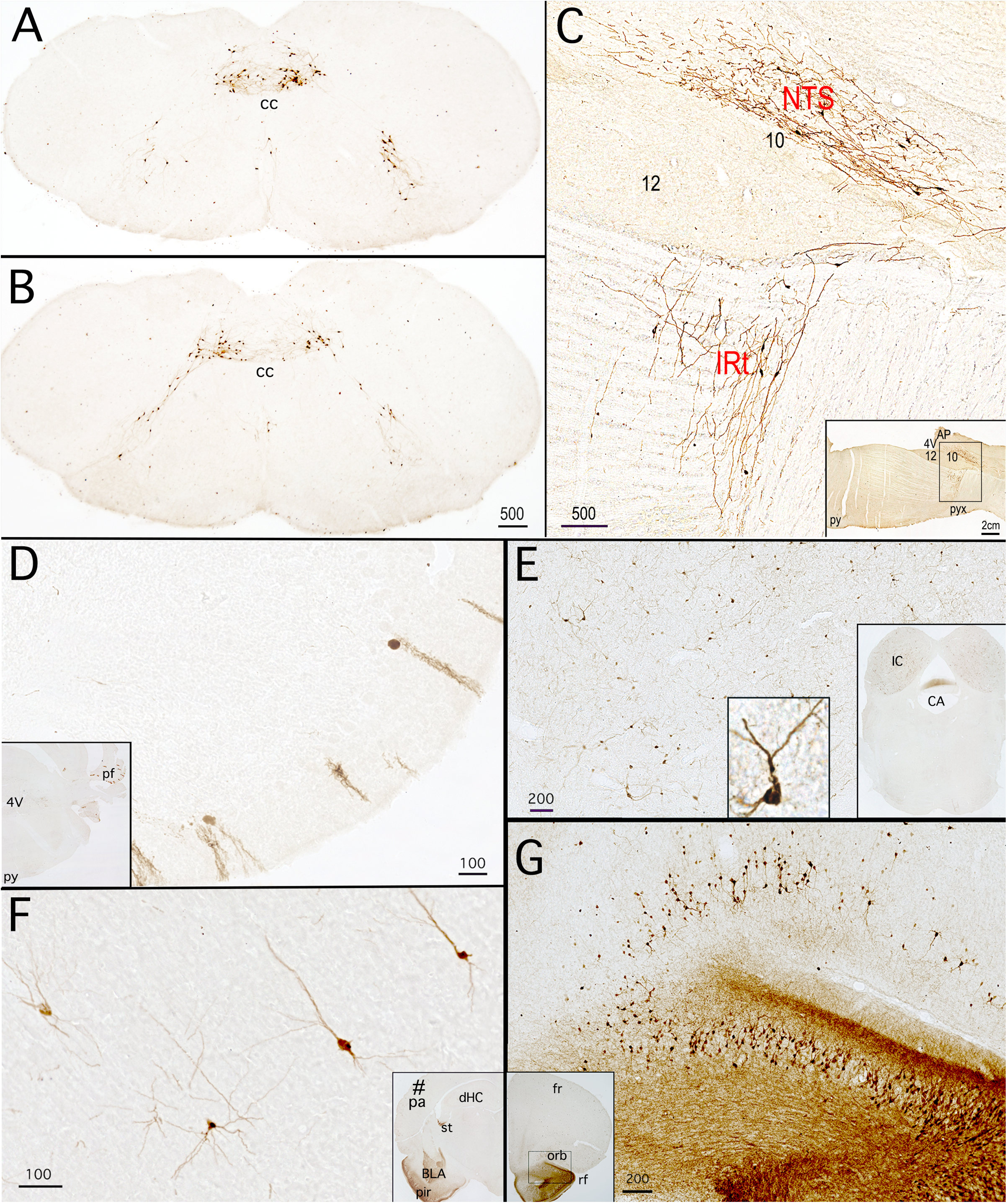
Central patterns of tdTom immunoperoxidase labeling in adult Gcg-Cre/tdTom reporter rats. **A, B**, coronal sections through two rostrocaudal levels of the cNTS and IRt. cc, central canal. **C**, parasaggital section through the caudal medulla, depicting tdTom-positive neurons and extensive dendritic branching in the cNTS and IRt. Inset shows the boxed region that is enlarged in panel C. *4V, fourth ventricle; 10, dorsal motor nucleus of the vagus; 12, hypoglossal nucleus; AP, area postrema; py, pyramidal tract; pyx, pyramidal decussation*. **D**, paraflocculus *(pf)* of cerebellum (shown at lower magnification in the inset). **E**, inferior colliculus (IC, shown at lower magnification in the inset). An example of a tdTom-positive neuron within the inferior colliculus is shown at higher magnification in the inset in lower center of panel E. *CA, cerebral aqueduct*. **F**, labeled pyramidal neurons within the piriform cortex *(pir)*. The inset in panel F shows a lower-magnification view of the piriform cortex and nearby basolateral amygdala *(BLA)*. See Figure 7 for additional images of BLA labeling. *dHC, dorsal hippocampus; st, stria terminalis*. **G**, ventral orbital *(orb)* and piriform cortex (shown at lower magnification in the inset). *fr, frontal cortex; rf, rhinal fissure*. Scale bars are in microns, except in the panel C inset.

#### 3.4.2 CNS reporter labeling in GLU-Cre/tdRFP mice

Reporter labeling in neonatal and adult Gcg-Cre/tdTom rats (described above, *3.4.1*) could result from transient developmental expression of *Gcg*, which would enable permanent cellular expression of *tdTom* after Cre-mediated excision of the floxed stop codon in the Rosa26 locus. To gain insight into this possibility, we compared the distribution of reporter labeling in Gcg-Cre/tdTom rats with reporter labeling in adult GLU-Cre/tdRFP mice, using tissue sections stored from a previous study [34]. Similar to reporter labeling in Gcg-Cre/tdTom rats, GLU (i.e., *Gcg*)-expressing cells in GLU-Cre/tdRFP mice are similarly marked in a permanent manner after Cre-mediated recombination of a floxed stop codon, such that transient developmental *Gcg* expression similarly enables permanent cellular reporter expression. These mice display highly specific hindbrain reporter labeling of GLP1 neurons within the cNTS and IRt [35]. Our new comparative analysis revealed that the more widespread distribution of persistent CNS reporter labeling observed in Gcg-Cre/tdTom rats (see *3.4.1*, **Fig. 6**) duplicates the distribution of reporter labeling in GLU-Cre/tdRFP mice, in which tdRFP-positive neurons are present in the same CNS regions (data not shown), including the BLA.

#### 3.4.3 Further examination of Gcg and iCre mRNA expression in the BLA

Given that robust tdTom reporter expression was observed within the BLA in both Gcg-Cre/tdTom rats and in GLU-Cre/tdRFP mice, we used RNAscope FISH to determine whether densely-packed collections of persistently reporter-labeled neurons in the BLA and adjacent piriform cortex express detectable levels of *Gcg* mRNA in neonatal and adult Gcg-Cre/tdTom rats. Scattered BLA (but not cortical) neurons contained detectable *Gcg* mRNA transcripts in neonatal (P10) rats (**Fig. S2D**), but not in adult rats (data not shown). However, the presence of functional iCre protein in a subset of reporter-labeled BLA neurons was demonstrated in adult Gcg-Cre/tdTom rats that received BLA-targeted microinjection of a Cre-dependent AAV expressing EGFP reporter (**Fig. S2A-C)**. EGFP labeled a subset of tdTom-positive BLA neurons within the AAV injection site, filling their spiny dendrites and axonal projections (**Fig. S2**).

#### 3.4.4 Pancreatic and intestinal reporter labeling in neonatal Gcg-Cre/tdTom rats

Reporter labeling in CLARITY-processed pancreatic and intestinal tissue from neonatal Gcg-Cre/tdTom rats (P2, P10) is illustrated in **Figure 7**. TdTom-positive cells were present in pancreatic islets and villi of the small intestine, and all tdTom reporter-labeled cells (100%) were GLP1-immunopositive.

**Figure 7.**
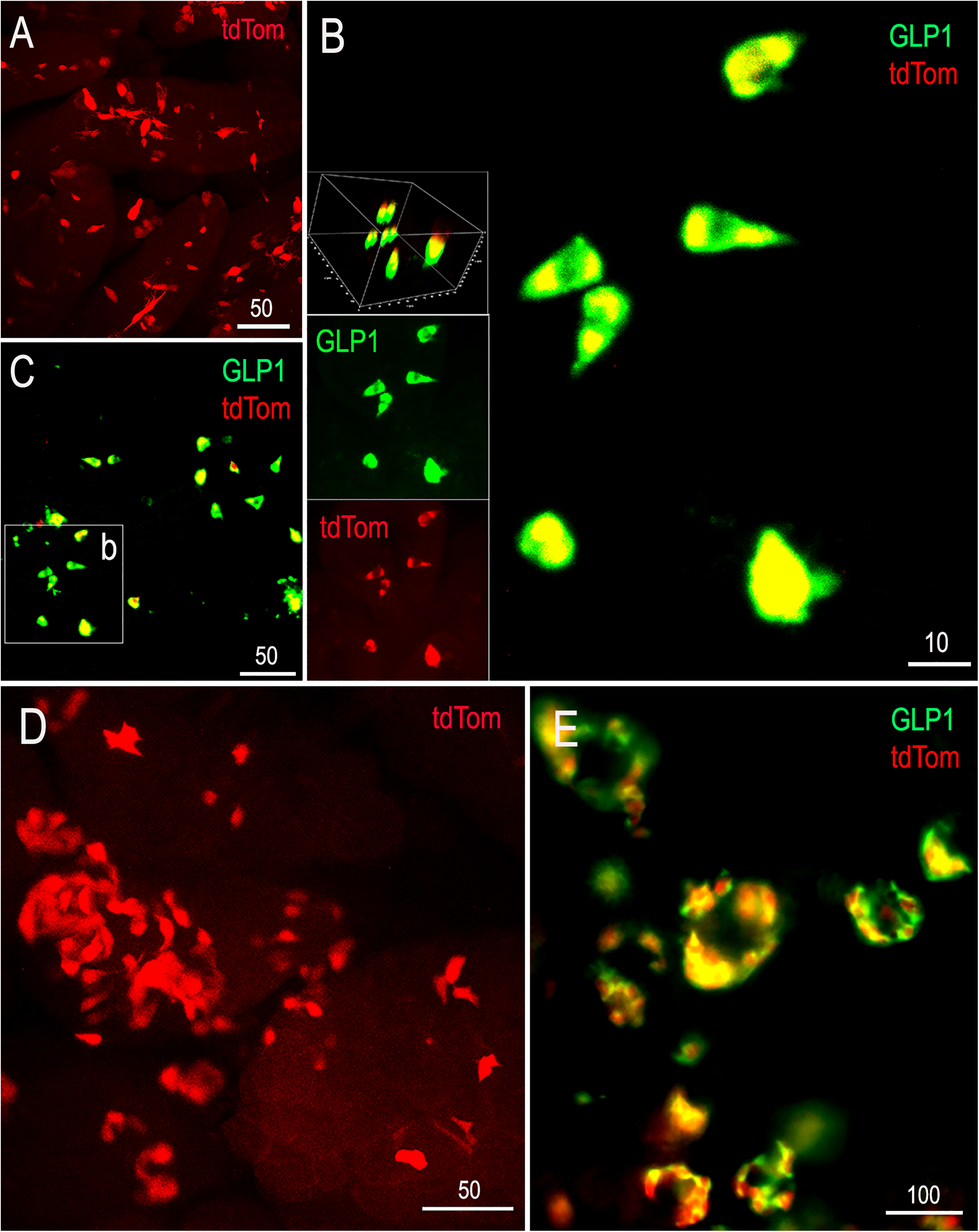
Pancreatic and intestinal tdTom native reporter (red) and GLP1 immunolabeling (green) in neonatal (P2, P10) Gcg-Cre/tdTom rats. Tissues were processed using CLARITY to remove lipid membranes and enhance visualization of fluorescent labeling. **A**, TdTom-positive cells within the intestinal villi (P10 rat). **B**, enlarged view of the boxed region (b) shown in panel **C** (P2 rat). Rotated views of confocally imaged tdTom-expressing intestinal cells demonstrate that all (100%) are GLP1-immunopositive. **D**, tdTom-labeled pancreatic cells (P10). **E**, immunolabeling for GLP1 demonstrates complete overlap with tdTom labeling in pancreatic islets (P2). Scale bars are in microns.

### 3.5 Cre-dependent anterograde tracing from Gcg-expressing cNTS neurons

AAVs expressing Cre-dependent EGFP or EYFP were microinjected into the cNTS in Homo, Het, and WT Gcg-Cre rats, followed by immunocytochemical enhancement of green reporter labeling to visualize reporter-labeled axonal projections. As expected, Cre-mediated reporter expression was observed within the cNTS injection site in Homo and Het Gcg-Cre rats, but not in WT rats born to Het Gcg-Cre dams/sires (**Fig. S3A-C**). Further, all (100%) EGFP/EYFP-positive cNTS neurons were double-labeled for GLP1 in Hets, and closely adjacent Dbh-positive A2 noradrenergic neurons were never reporter-labeled in Het or Homo rats (**Fig. S3D**). The axonal projections of EGFP/EYFP-labeled, *Gcg*-expressing cNTS neurons were visualized in all subcortical CNS regions that have been reported to contain GLP1-positive axons in adult rats and mice, including the medulla, pons, midbrain, diencephalon, and limbic forebrain [36][37][38]. **Figure S3E** shows reporter-labeled axons within the paraventricular nucleus of the hypothalamus that arise from transfected cNTS neurons in a representative Homo Gcg-Cre rat.

### 3.6 Confirmation of chemogenetic activation of Gcg-expressing cNTS neurons

In a pilot study, female Het Gcg-Cre rats received bilateral cNTS microinjection of AAV8 expressing Cre-dependent mCherry and GqDREADD, or a control AAV expressing only the mCherry reporter. Rats were later injected i.p. with either 0.15M saline vehicle, or vehicle containing CNO (1.0mg/kg BW) before perfusion. Brainstem sections were processed for dual immunocytochemical labeling of nuclear cFos and cytoplasmic mCherry reporter (**Fig. S4**). ANOVA revealed significant main effects of AAV [F(1,14) = 6.08, *P* = 0.027] and i.p. injection [F(1,14) = 26.0, *P* = 0.0002] on the proportion of mCherry-labeled neurons that expressed cFos, as well as a significant interaction between AAV and i.p. injection conditions [F(1,14) = 33.78, *P* < 0.0001]. Compared to activation after i.p. saline, CNO did not increase cFos activation in rats with control mCherry reporter virus expression (*P* = 0.86; **Fig. S4E,F**; *CTRL*). Conversely, CNO produced a 15-fold increase in the proportion of mCherry-positive, GqDREADD-expressing neurons activated to express cFos (*P* < 0.0001; **Fig. S4F**, *Gq*).

### 3.7 Chemogenetic activation of Gcg-Cre neurons reduces food intake

#### 3.7.1 Intra-cNTS administration of CNO

In the first food intake test, rats had a choice between consuming chow and peanut butter. In WT control rats (which do not express *iCre*) that earlier received cNTS-targeted injection of AAV expressing Cre-dependent GqDREADD, intra-cNTS delivery of CNO did not alter chow or peanut butter intake compared to intra-cNTS vehicle (**Fig. S5)**, and postmortem examination of hindbrain sections revealed no fluorescent AAV reporter labeling (not shown). Conversely, in Het Gcg-Cre rats, separate ANOVAs at the 1 and 16h timepoints revealed a significant main effect of sex [F(1,13) = 15.83, *P* = 0.002] and a significant interaction between sex and CNO treatment on 16h chow intake [F(1,13) = 10.65, *P* = 0.006], but no main effect of CNO at 16h [F(1,13) = 2.487, *P* = 0.13]. In male rats, chow intake was suppressed after intra-cNTS CNO injection at the 16h timepoint, but not at the 1h timepoint (**Fig. S5G,H**). There was no main effect of CNO on chow intake in female rats, although there was a strong trend (*P* = 0.05) for CNO to reduce peanut butter intake in females at the 1h timepoint [CNO x sex, F(1, 13) = 9.426, *P* = 0.008; CNO F(1, 13) = 0.3, *P* = 0.5903; sex F(1,13) = 0.06728, *P* = 0.8]. CNO did not alter peanut butter intake in males at 1h, and did not alter peanut butter intake at 16h in either sex (**Fig. S5I,J)**.

#### 3.7.2 Central Exendin-9 (Ex9) blockade of the hypophagic effect of central CNO in male rats

Male Gcg-Cre Het rats (N=8) that were used for the chow-peanut butter choice experiment (*3.7.1*, above) were used in a follow-up experiment to determine whether the ability of central (in this experiment, LV-delivered) CNO to suppress chow intake depends on central GLP1R signaling. As shown in **Figure 8B**, male rats reduced their 24h chow intake after LV delivery of CNO, and the hypophagic effect of CNO at 24h was completely blocked by prior LV administration of Ex9 (**Fig. 8B)** [CNO x EX9 F(1,14) = 11.39, *P* = 0.005; CNO F(1,14) = 1.169, *P* = 0.3; EX9 F(1,14) = 8.433, *P* = 0.02]. As expected given these intake data, CNO reduced body weight at the 24h timepoint in rats pretreated with vehicle, whereas rats pretreated with EX9 before CNO did not lose weight (**Fig. 8C)** [CNO x EX9 F(1,14) = 4.259, *P* = 0.05; CNO F(1,14) = 4.428, *P* = 0.05; EX9 F(1, 14) = 8.433, *P* = 0.22]. At the 1h timepoint, CNO treatment tended to reduce intake in rats pretreated with vehicle, although this effect did not reach significance (**Fig. 8A)** [CNO x EX9 F(1, 11) = 1.033, *P* = 0.3; CNO F(1, 14) = 3.688, *P* = 0.07; EX9 F(1,11) = 2.181, *P* = 0.16]. Three data points are absent from the 1h food intake analysis, two due to spillage and one determined to be an outlier as indicated by Grubbs test.

**Figure 8.**
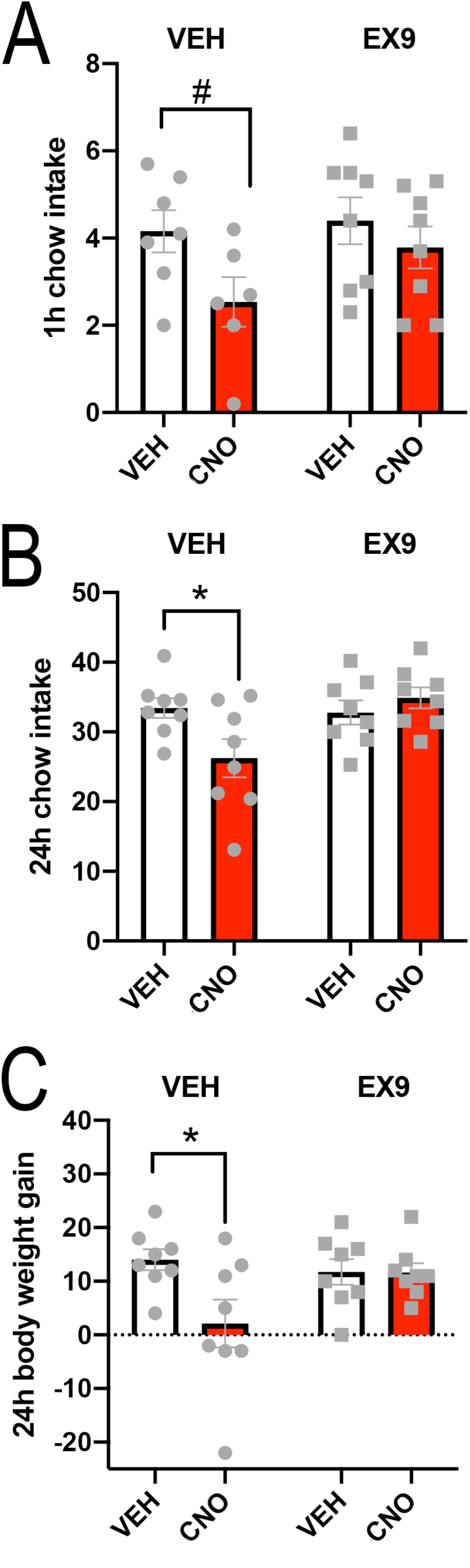
Central Exendin-9 (Ex9; a GLP1R antagonist) blockade of the hypophagic effect of central CNO. Male Gcg-Cre Het rats with prior cNTS-targeted delivery of AAV expressing Cre-dependent GqDREADD were used to determine whether central (LV-delivered) CNO suppresses dark-onset chow intake, and if so, whether this effect depends on central GLP1R signaling. **A**, 1h chow intake was not significantly reduced by LV administration of vehicle followed by either LV vehicle or LV CNO, although a trend for CNO to suppress 1h intake was evident. **B**, 24h chow intake was significantly reduced when LV vehicle was followed by LV delivery of CNO (\**P* < 0.05), and this 24h hypophagic effect was completely blocked by LV delivery of CNO was preceded by LV delivery of Ex9. **C**, CNO reduced body weight gain at the 24h timepoint in rats pretreated with vehicle, whereas rats pretreated with EX9 before CNO did not lose weight.

#### 3.7.3. Systemic administration of CNO

A separate cohort of male Het Gcg-Cre rats with bilateral cNTS-targeted injections of Gq DREADD virus (AAV8) was housed in BioDaq cages for automated dark-onset overnight food intake measurements after i.p. injections of 0.15M NaCl vehicle, vehicle containing CNO (1.0mg/kg BW), and/or vehicle containing a low dose of CCK (1.0μg/kg BW). Within-subjects repeated measures ANOVA revealed significant main effects of i.p. treatment condition [F(2.6,34) = 9.2; *P* = 0.0003] on overnight chow intake, and a significant interaction between i.p. treatment and time [F(3.8,49) = 3.0; *P* = 0.03]. Average food intake over the 12h dark cycle in Gcg-Cre rats after each of the four i.p. treatment combinations is plotted in **Figure 9A**. Compared to overnight food intake after control treatment (i.e., saline vehicle followed by saline vehicle; saline-saline), post-hoc t-tests corrected for multiple comparisons revealed that CNO-saline did not inhibit food intake at dark hour (D1), D2, D4, or D6. However, a delayed cumulative hypophagic effect of CNO-saline reached statistical significance by D12 (*P* = 0.045). Saline-CCK did not suppress food intake at any timepoint compared to intake after saline-saline. However, when CNO was followed by CCK (i.e., CNO-CCK), a significant hypophagic effect was evident at D4 (*P* = 0.022), D6 (*P* = 0.0008), and D12 (*P* = 0.017). **Figure 9B** shows individual within-subjects data points for cumulative intake by each rat after each i.p. injection condition at D4, D6, and D12, with intake expressed as % reduction compared to intake by the same rat after control (saline-saline) treatment. ANOVA revealed a significant main effect of i.p. treatment [F(1.8,23) = 9.0; *P* = 0.0018] and a significant interaction between i.p. treatment and time [F(2.9,37) = 3.6; *P* = 0.025]. Post-hoc tests corrected for multiple comparisons revealed that CNO-CCK suppressed food intake to a greater degree than CCK alone at D4 (*P* = 0.013) and D6 (*P* = 0.011), and suppressed intake more than CNO alone at D6 (*P* = 0.012).

**Figure 9.**
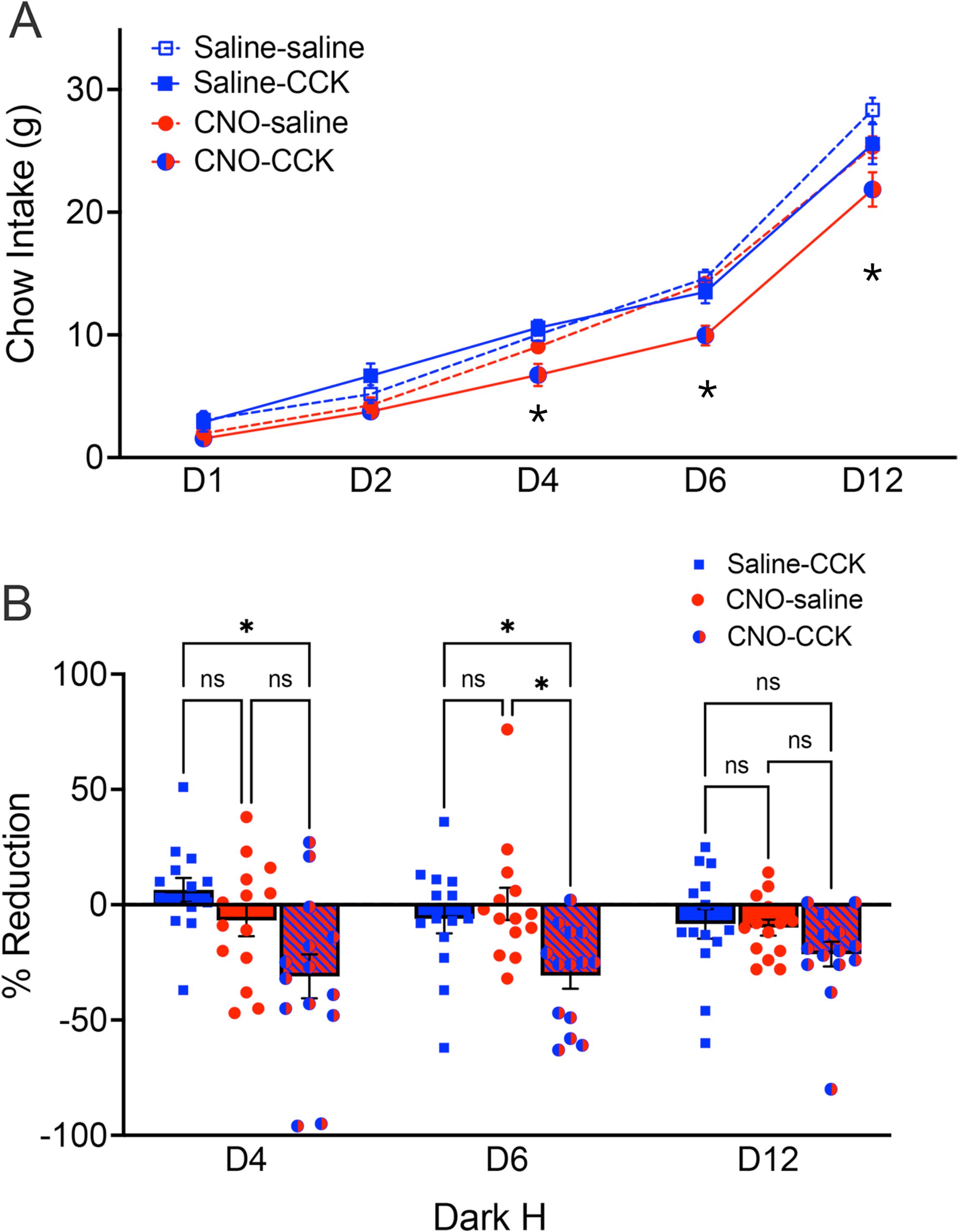
Overnight chow intake in male Het Gcg-Cre rats (N=14) with bilateral cNTS-targeted injections of Gq DREADD virus after i.p. injections of saline followed by saline (control, saline-saline), saline followed by CCK (saline-CCK), CNO followed by saline (CCK-saline), or CNO followed by CCK (CNO-CCK). **A**, average food intake over the 12h dark cycle grouped by i.p. treatment combination. Compared to overnight food intake after saline-saline treatment, CNO-saline appeared to inhibit food intake after dark hour (D)4, although the cumulative hypophagic effect of CNO-saline did not reach significance until D12 (\**P* = 0.045 vs. saline-saline). Saline-CCK did not suppress food intake at any timepoint compared to intake after saline-saline. However, a significant hypophagic effect of CNO-CCK was evident at D4 (*P* = 0.022), D6 (*P* = 0.0008), and D12 (*P* = 0.017). **B**, individual within-subjects data for cumulative intake by rats after each i.p. injection condition at D4, D6, and D12, with intake expressed as % reduction compared to intake by the same rat after control (saline-saline) treatment. CNO-CCK suppressed food intake to a greater degree than CCK alone at D4 (\**P* = 0.013) and D6 (\**P* = 0.011), and suppressed intake more than CNO alone at D6 (\**P* = 0.012).

## 4.0 Discussion

Our results demonstrate the utility of a new Gcg-Cre rat model to interrogate the anatomy and function of *Gcg*-expressing cells in the brain and periphery. Here we report evidence that Het (+/-) and Homo (+/+) Gcg-Cre rats breed and develop normally, with Het and Homo rats displaying relatively minor genotype- and sex-dependent differences in body weight and fat mass when maintained on chow. We document a high selectivity of hindbrain, pancreatic, and intestinal cellular co-expression of *Gcg, iCre*, and/or GLP1 immunolabeling. We also provide new evidence that *Gcg* is transiently expressed in additional brainstem and forebrain regions during development in rats and in mice, and present evidence that a low level of *Gcg* promoter-driven *iCre* expression persists within the adult BLA. We also report our unexpected finding that insertion of an IRES for *iCre* expression near the 3’ UTR of *Gcg* markedly reduces *Gcg* mRNA and GLP1 protein levels, such that Homo Gcg-Cre rats present a unique *Gcg* knockdown/knockout rat model; conversely, Het Gcg-Cre rats have normal brain GLP1 levels and moderately reduced plasma levels. Finally, by using Het Gcg-Cre rats, we demonstrate that chemogenetic excitation of Gcg-expressing cNTS neurons reduces dark-onset food intake in a sex- and central GLP1R-dependent manner, and amplifies the vagally-mediated hypophagic effect of systemic CCK.

### 4.1 General health of Gcg-Cre rats

Compared to WT Sprague-Dawley rats, Het and Homo Gcg-Cre rats born in the FSU colony and at the Janvier facility displayed normal breeding competence, survival rates, growth curves, and overall health. Genotype had no effect on BW in males, although female Homo rats tended to weigh slightly more than age-matched Het and WT rats. Conversely, genotype did not impact body fat mass in females, whereas Homo males had less body fat than Hets. It should be noted that these parametric data were collected from rats maintained on normal rat chow. The potential impact of developmental and adult dietary conditions on body energy homeostasis and fat mass in Gcg-Cre rats is certainly of interest.

### 4.2 Plasma glucose levels in Gcg-Cre rats

Fed and fasted plasma glucose levels were similar in WT and Het Gcg-Cre rats, and comparable to levels reported previously in WT SD rats [39]. However, Homo Gcg-Cre rats of both sexes had significantly lower plasma glucose levels than WT and Het rats under fed conditions, and glucose levels fell in WT and Het rats but not in Homo rats after overnight food deprivation. Intestinal enteroendocrine and pancreatic islet cells serve as sources of glucagon, GLP1, and other *Gcg*-derived peptides with glucoregulatory action, and fasting increases pancreatic glucagon production [40]. Whole-body *Gcg* knockout mice exhibit resistance to the development of glucose intolerance due to their deficit in pancreatic glucagon, which normally stimulates hepatic gluconeogenesis [16]. We did not assess plasma glucagon levels or *Gcg* expression in the pancreas or intestine in the present study, but reduced pancreatic glucagon production in Homo Gcg-Cre rats could explain their lower plasma glucose levels under fed conditions. However, it is more difficult to explain the absence of fasting-induced hypoglycemia in Homo Gcg-Cre rats. The apparent discrepancy in the effect of whole-body protein knockdown/knockout to reduce basal glucose levels in rats (present study) vs. acute stimulation of hindbrain *Gcg* neurons to reduce glucose production in mice [19] could reflect a species difference, or could point towards different roles of peripheral vs. central *Gcg*-encoded proteins in glucose homeostasis. Additional research will be necessary to understand why overnight fasting did not further reduce plasma glucose levels in Homo rats. Transgenic mice with gut-specific knockdown of *Gcg* expression display markedly reduced circulating levels of active GLP1, but normal plasma insulin levels and normal i.p. glucose tolerance [15]. In that report, fasting glucose levels also were normal in mice with gut-specific *Gcg* knockdown; however, glucose levels under fed conditions were not reported, and so the effect of fasting was not determined [15].

### 4.2 Specificity of iCre expression in Gcg-Cre rats

*Gcg* expression within the rat brain exhibits region- and developmental stage-specific differences, with levels often higher during early development. For example, *Gcg* mRNA transcripts detected using qRT-PCR in the developing rat cortex, hippocampus, and cerebellum at embryonic day 19 decline to very low levels in the adult, whereas brainstem *Gcg* mRNA expression relatively constant after embryonic day 19 [41]. In the present study, cellular patterns of tdTom reporter labeling appeared identical in neonatal and adult Gcg-Cre/tdTom rats. This is unsurprising, because reporter labeling should identify cells within the brain and periphery that express *Gcg* at any point from gestation, even if cellular expression is developmentally transient (e.g., see [42]). In cleared intestinal and pancreatic tissue samples from neonatal rats, 100% of tdTom reporter-labeled cells were immunopositive for GLP1. Although pancreatic protein levels of *Gcg*-encoded glucagon are much higher, GLP1 is a known cleavage product in pancreatic α-cells [15][43]. Reporter-labeled intestinal cells displayed a distribution and morphology consistent with enteroendocrine L-cells [44], while reporter-labeled pancreatic cells displayed a distribution and morphology consistent with α-cells [45].

Within the hindbrain cNTS, IRt, and midline raphe, tdTom reporter labeling displayed high efficacy and specificity, being 100% colocalized with GLP1 immunolabeling and/or *Gcg* mRNA transcript labeling assessed using RNAscope. However, tdTom reporter labeling was not restricted to the hindbrain, but included several higher brain regions reported to lack detectable *Gcg* expression in adults. In addition, the olfactory bulb in Gcg-Cre/tdTom rats contained many more reporter-labeled cells than the small population of interneurons that *Gcg* in adult mice and rats [7][33][6], and in the present study only a subset of tdTom-labeled olfactory bulb cells contained RNAscope-detectable *Gcg* mRNA. Particularly robust tdTom reporter labeling was observed within the BLA and piriform cortex in neonatal and adult Gcg-Cre/tdTom rats, but RNAscope revealed no above-background levels of *Gcg* mRNA expression in adults.

We compared the pattern of central tdTom labeling in our rats with reporter labeling in adult GLU-Cre/tdRFP mice, in which GLU- (i.e., *Gcg)*-expressing cells are similarly marked in a permanent manner after Cre-mediated recombination of a floxed stop codon [46][47]. In these mice, tdRFP-immunolabeled neurons were identified in the same hindbrain, midbrain, diencephalic and telencephalic brain regions that contained reporter labeling in Gcg-Cre/tdTom rats. To our knowledge, the detailed CNS distribution of reporter-labeled cells in adult GLU-Cre/tdRFP mice has not previously been reported. Using a different Gcg-Cre mouse line crossbred to floxed tdTom reporter mice, Gaykema and colleagues described “very limited off-target expression (1-2 neurons per 40-μm section)” (*data not shown [25])* within most of the brain regions that we observed to contain large numbers of reporter-labeled cells in Gcg-Cre/tdTom rats and in GLU-Cre/tdRFP mice. Conversely, cellular reporter labeling in PPG-YFP mice, which depends on active *Gcg* transcription, has been reported to be anatomically restricted to the cNTS, IRt, and midline raphe, with scattered neurons present within the piriform cortex, but not within the BLA [36].

Apart from the olfactory bulb, *Gcg* mRNA expression has not been reported within the telencephalon in adult rodents, and (as mentioned above) *Gcg* mRNA within the BLA and piriform cortex was not detectable using RNAscope in adult Gcg-Cre/tdTom reporter rats. However, images available online through the Allen Brain Atlas [48] depict a small population of BLA (but not piriform cortex) neurons with low to moderate levels of *Gcg* expression in adult mice (https://mouse.brain-map.org/experiment/show/75749416). Those data fit with our present results indicating that a subset of tdTom-positive BLA neurons were transfected in adult Gcg-Cre/tdTom rats after BLA-targeted microinjection of AAV expressing a Cre-dependent green fluorescent reporter. Microinjection of the same AAV into the cNTS in WT Gcg-Cre rats did not result in labeling, evidence that observed BLA labeling in Gcg-Cre rats is Cre-dependent. Since cellular *iCre* expression depends on active *Gcg* transcription, and since extremely low levels of Cre-recombinase are sufficient for expression of Cre-dependent AAV products, we interpret these results as evidence that *Gcg* continues to be expressed at low levels (i.e., undetectable using RNAscope FISH) by a subset of adult BLA neurons. Additional work will be needed to evaluate potential iCre expression in the piriform cortex or other reporter-labeled brain regions that lack detectable *Gcg* expression and GLP1-immunopositive neurons, and to evaluate whether low basal levels of *Gcg* expression might increase in response to physiological or experimental conditions. The potential for Cre-mediated transcription in unexpected and/or undesired regions should always be evaluated when using Cre-driver mice and rats for targeted manipulation of specific cells.

### 4.3 Disruption of Gcg mRNA expression and GLP1 protein in Homo Gcg-Cre rats

In order to generate rats with Cre-recombinase directed to the glucagon gene (*Gcg*), an IRES for *iCre* was inserted after the final coding sequence of exon 6 in the *Gcg* genetic sequence, before the 3′ UTR. While this strategy was designed to retain endogenous *Gcg* expression, qRT-PCR and quantitative RNAscope-based FISH revealed a marked reduction of *Gcg* mRNA transcript levels within the cNTS in Homo Gcg-Cre rats, and a moderate reduction in Hets when measured by RNAscope (but not by qRT-PCR). qRT-PCR confirmed highly correlated expression of *iCre* and *Gcg* within the cNTS in Het rats, whereas *Gcg* expression in Homo rats was almost non-detectable despite *iCre* expression levels similar to those in Hets. Since RNA sequences in 3′ UTRs serve to regulate mRNA stability [49], *Gcg* mRNA stability could be disrupted in our model, leading to lower levels of GLP1 and other *Gcg*-encoded proteins. Alternatively, or in addition, insertion of the IRES-iCre cassette may have affected splicing of the *Gcg* gene. If so, the pre-mRNA transcript containing introns would be generated as expected, with the *iCre* sequence translated from the IRES site; however, *Gcg* mRNA wouldn’t be spliced properly, leading to its degradation. This could explain why *iCre* expression is comparable in Het and Homo Gcg-Cre rats, whereas *Gcg* expression is almost completely disrupted in Homo rats. Similar disruption of target genes and their corresponding proteins has been reported in other IRES-Cre models [e.g. [50][51]]. It is important to be aware of such disruption when it occurs, as it can affect experimental results and their interpretation; however, potential disruption is often unexamined (or unreported) in published studies using Cre-driver model organisms.

As expected, given the near-absence of detectable *Gcg* mRNA within the cNTS in Homo Gcg-Cre rats, GLP1 protein levels were correspondingly and markedly reduced in the brainstem, forebrain, and plasma. In Het rats with disrupted *Gcg* expression from only one genomic allele, average GLP1 protein levels were reduced by approximately 50% in the plasma, but were not reduced within the brain. We did not assess *Gcg* mRNA expression in peripheral tissues, but genotype-related effects likely parallel those in the brain. We predict markedly less *Gcg* expression in the pancreas and intestine of Homo rats but only a partial reduction in Hets. Our finding that GLP1 protein levels in Hets are reduced in plasma but not brain is likely related to the rapid degradation of peripherally released GLP1 by blood-borne dipeptidyl peptidase 4 [52][53][54], whereas GLP1 produced centrally and transported axonally by neurons within the cNTS and IRt is protected within dense core vesicles.

### 4.4 Widespread axonal projections of Gcg-expressing cNTS neurons

Cre-dependent AAVs expressing an anterogradely-transported fluorescent reporter were microinjected into the cNTS in Homo, Het, and WT Gcg-Cre rats, followed by immunocytochemical enhancement of reporter labeling to visualize the axonal projections of transfected neurons. As expected, no cellular or axonal labeling was observed in WT rats, which lack Cre expression. In Het rats, all labeled neurons within the cNTS injection site were GLP1-positive. In Homo rats, some reporter-labeled neurons did not express detectable GLP1 immunolabeling, likely due to disruption of *Gcg* expression (discussed above, *4.3*). However, noradrenergic A2 neurons within the cNTS were never transfected in either Het or Homo rats, despite their intermingled distribution among GLP1 neurons. Reporter-labeled axons were observed in multiple brainstem, hypothalamic, and limbic forebrain regions, fully consistent with previous reports mapping the widespread distribution of GLP1-positive fibers arising from the hindbrain in adult rats and mice [36][37][38]. We only injected tracing AAVs into the cNTS in this experiment, and so additional work will be needed to determine whether the axonal projections of *Gcg*-expressing neurons in the cNTS differ from those of *Gcg*-expressing neurons in the IRt or midline raphe region.

### 4.5 Chemogenetic stimulation of Gcg-expressing cNTS neurons suppresses food intake in a sex-dependent manner, and depends on central GLP1R signaling

Published work using Cre-driver mice indicates that chemogenetic activation of cNTS GLP1 neurons is sufficient to suppress food intake [11][25][34]. The present study sought to confirm and extend those findings using Gcg-Cre rats. For this purpose, AAV2 or AAV8 constructs expressing the same Cre-dependent excitatory DREADD were delivered to the cNTS. Experimental controls included administration of CNO to non-DREADD-expressing WT rats to control for off-target activity [55]. We also used a crossover design to compare food intake within the same subjects after CNO vs. vehicle injection [56].

In the first chemogenetic experiment, adult male and female Het Gcg-Cre rats with AAV2-mediated DREADD expression subsequently received unilateral parenchymal cNTS delivery of vehicle or vehicle containing CNO, followed by a dark-onset food intake test in which rats were presented with both chow and peanut butter. Overnight (16h) chow intake was reduced in male rats after CNO treatment, but CNO did not suppress peanut butter intake. Conversely, CNO did not suppress chow intake in female rats, although there was a trending effect for CNO to reduce 1h peanut butter intake in females. A follow-up experiment demonstrated that the ability of LV-delivered CNO to suppress overnight chow intake in male rats depended on central GLP1R signaling, as the hypophagic effect was blocked by prior LV administration of Ex9.

In the second experiment, male Het Gcg-Cre rats with bilateral AAV8-mediated DREADD expression within the cNTS received a low systemic dose of CNO, either alone or followed by a behaviorally subthreshold dose of CCK. CCK administration was used to selectively activate gastrointestinal vagal afferents that express CCK receptors and normally contribute to meal-induced satiety [57]. CNO delivered i.p. led to a small but significant suppression of overnight chow intake, whereas CCK by itself did not. When CNO was followed by CCK in the same animal, the hypophagic effect of CNO was amplified and appeared earlier during the dark cycle. Thus, chemogenetic activation of GLP1 neurons within the cNTS appears to amplify vagal sensory signaling induced by systemic CCK, thereby enhancing CCK-induced hypophagia.

The combined results of these food intake experiments support the view that chemogenetic activation of GLP1 neurons within the cNTS can suppress food intake in rats, as previously demonstrated in mice. The effect in rats appears to be sex-dependent, with male rats more sensitive than females. It is interesting that CNO by itself was less effective to suppress food intake in the second experiment in which CNO was delivered systemically, since overnight chow intake was suppressed by approximately 25% in male rats after unilateral cNTS-targeted delivery of CNO. This difference in results could be related to different AAV serotypes for the GqDREADD used in each experiment (AAV2 vs. AAV8), and/or to different CNO delivery routes (cNTS vs. i.p.) that would affect stimulation of GqDREADD expressed unilaterally or bilaterally by GLP1 neurons within the cNTS.

## 5.0 Conclusions

Gcg-Cre rats are a valuable addition to the research toolbox for examining the structure and function of central neurons and peripheral secretory cells that express *Gcg*. Our discovery that Homo Gcg-Cre rats have markedly reduced *Gcg* mRNA and GLP1 protein levels make them a useful and novel rodent model to address research questions regarding the role of GLP1 and other *Gcg*-encoded protein products in food intake, metabolism, obesity, and diabetes. It will be important to use Het Gcg-Cre rats in experiments that assume or rely upon normal-range levels of endogenous GLP1 and other *Gcg*-encoded proteins.

## SUPPLEMENTARY FIGURES

**Supplementary Figure 1**. Body weight (BW) in young Gcg-Cre rats bred in the Janvier facility. ***Left***, Male Gcg-Cre rats display similar growth curves and BW from weeks 6-9 postnatal, regardless of genotype. ***Right***, Female Homo Gcg-Cre rats display slightly higher BW compared to WT rats at weeks 8 and 9, but the difference was not significant (^#^*P* < 0.1 and > 0.05).

**Supplementary Figure 2**. Panels A-C2 depict epifluorescent (A) and confocal (B, C1, C2) images of EGFP viral labeling (green) in an adult female Gcg-Cre/tdTom reporter rat after microinjection of cre-dependent AAV expressing EGFP reporter. **A**, BLA injection site (b, boxed inset shown at higher magnification in panel B). Red (RFP) labeling is immunofluorescently enhanced tdTom reporter labeling. Blue DAPI counterstain. **B**, EGFP-expressing BLA neurons give rise to extensive dendritic and axonal processes. In the boxed inset (c), white arrows point out two neurons that are double-labeled for RFP (tdTom reporter) and EGFP. Boxed inset is shown at higher magnification in **C1** (RFP channel) and **C2** (EGFP channel). Many RFP-positive neurons do not express EGFP viral reporter. **D**, RNAscope FISH reveals Gcg mRNA expression in a subset of BLA tdTom (RFP)-positive neurons in a neonatal (P10) Gcg-Cre/tdTom reporter rat. White arrows point out double-labeled cells. Scale bar in B is in microns.

**Supplementary Figure 3**. Cre-dependent EGFP reporter labeling in adult Homo (**A**, +/+), Het (**B**, -/-), and WT (**C**, -/-) Gcg-Cre rats after cNTS-targeted AAV injections. No reporter labeling is observed in WT rats, which lack *iCre* expression. *cc, central canal*. **D**, inset shows boxed region enlarged in panel D, from a Het Gcg-Cre rat. All EGFP-positive cells are GLP1-positive, and vice versa, as shown in the 3 smaller panels below D. DBH-positive noradrenergic neurons are not transfected by the Cre-dependent AAV. **E**, EGFP-positive axons within the paraventricular nucleus of the hypothalamus in a Homo rat that received cNTS-targeted AAV injection. *V, 3*^*rd*^ *ventricle*. Scale bars are in microns.

**Supplementary Figure 4**. Confirmation of GqDREADD expression and CNO-induced activation of cNTS neurons in adult female Het Gcg-Cre rats. Tissue sections were double-labeled for immunoperoxidase localization of nuclear cFos (blue/black) and mCherry cytoplasmic reporter (brown). Rats received cNTS-targeted injections of Cre-dependent AAV expressing GqDREADD (Gq) or control AAV (CTRL) expressing only mCherry. Rats were injected i.p. with either saline vehicle or CNO (1 mg/kg BW) 90 min before perfusion. **A**, GqDREADD-expressing cNTS neurons are not activated to express cFos after i.p. saline. **B**, GqDREADD-expressing cNTS neurons are activated after i.p. injection of CNO. **C**, Control virus mCherry-labeled cells are not activated to express cFos after i.p. saline injection. **D**, Control virus mCherry-labeled cells are not activated to express cFos after i.p. CNO. **E**, summary data quantifying the proportion of mCherry-positive transfected neurons expressing cFos in rats injected with control virus (CTRL) or with GqDREADD-expressing virus (Gq). CNO does not increase activation in CTRL rats, but markedly increases activation in Gq rats (\*\**P* < 0.001). **F**, summary data quantifying the proportion of cFos-positive cNTS neurons that are mCherry labeled. CNO does not increase this proportion in CTRL rats, but significantly increases this proportion in Gq rats (\**P* < 0.05).

**Supplementary Figure 5**. *Left*, data from wildtype control (-/-) rats. Chow and peanut butter intake (**A-D**, 2-food choice test, **E**, chow only) in male and female WT Gcg-Cre rats with cNTS AAV injections of Cre-dependent GqDREADDs, followed by intra-cNTS administration of CNO (red bars) or vehicle (open bars) before dark-onset food access. **A, B**, 1h intake of chow (A) or peanut butter (B) does not differ in either sex following vehicle vs. CNO treatment. **C, D**, 16h overnight intake of chow (C) or peanut butter (D) does not differ in either sex following vehicle vs. CNO treatment. **E**, 24h chow intake (no choice) does not differ in male rats after vehicle vs. CNO treatment. **F**, body weight gain does not differ in male rats after vehicle vs. CNO treatment. Data from Het Gcg-Cre rats. Chow vs. peanut butter intake (2-food choice test) in male and female Het Gcg-Cre rats with cNTS AAV injections of Cre-dependent GqDREADDs, followed by intra-cNTS administration of vehicle (open bars) or CNO (red bars) before dark-onset food access. *Right*, data from Het (+/-) Gcg-Cre rats. **G**, 1h dark-onset chow intake; **H**, 16h overnight chow intake; **I**, 1h dark-onset peanut butter intake; **J**, 16h overnight peanut butter intake. At the 1h timepoint, CNO did not inhibit either chow (A) or peanut butter (C) intake in either sex compared to intake after vehicle treatment, although a strong trend towards inhibition of peanut butter intake was evident in females (panel C, ^#^*P* = 0.05). At the 16h timepoint, CNO significantly inhibited chow intake in male rats (panel B, \*\**P* < 0.05) but not in females. Peanut butter intake at 16 hr (D) did not differ in either sex after vehicle vs. CNO treatment.

