## Supplementary figures and images for "A Cre-Driver Rat Model for Anatomical and Functional Analysis of Glucagon *(Gcg)*-Expressing Cells in the Brain and Periphery"

### Supp Fig. 1

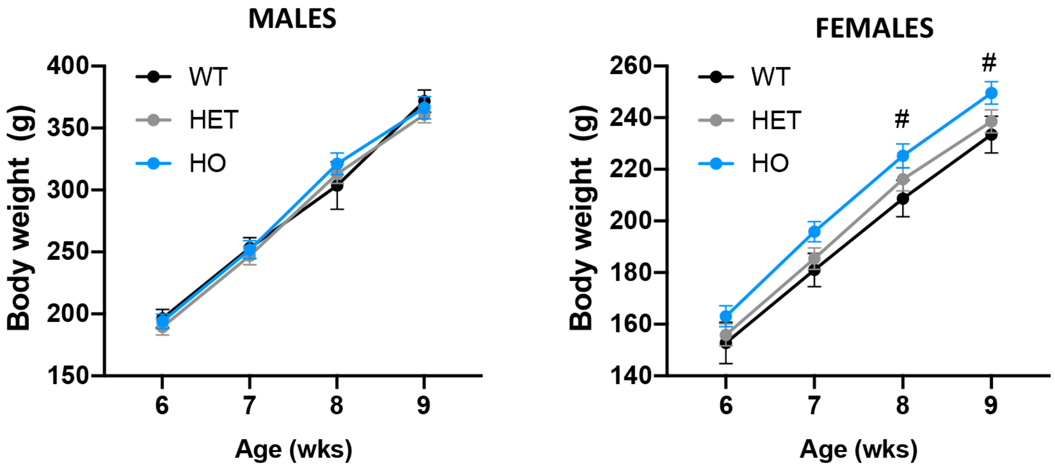

### Supp Fig. 2

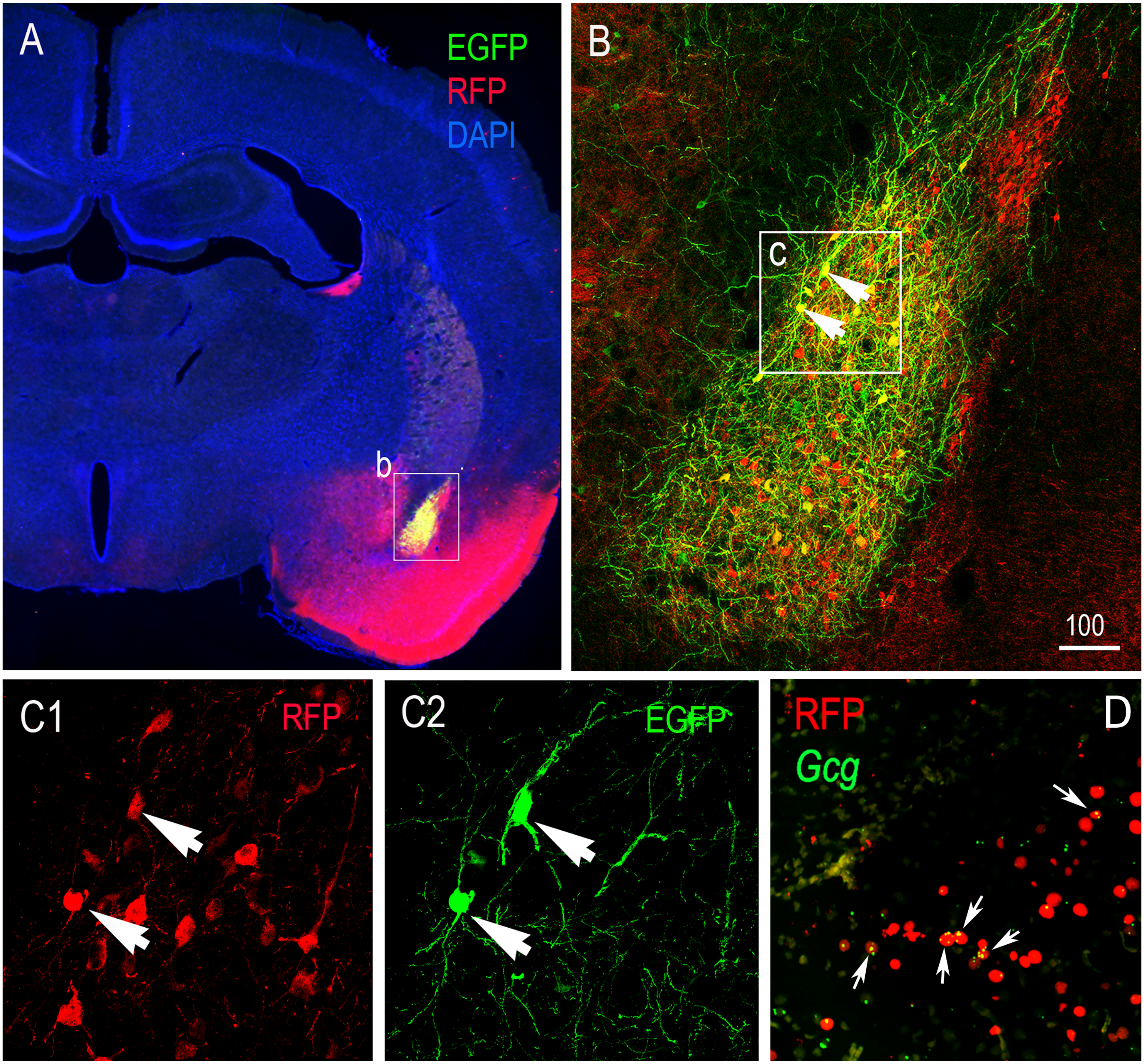

### Supp Fig. 3

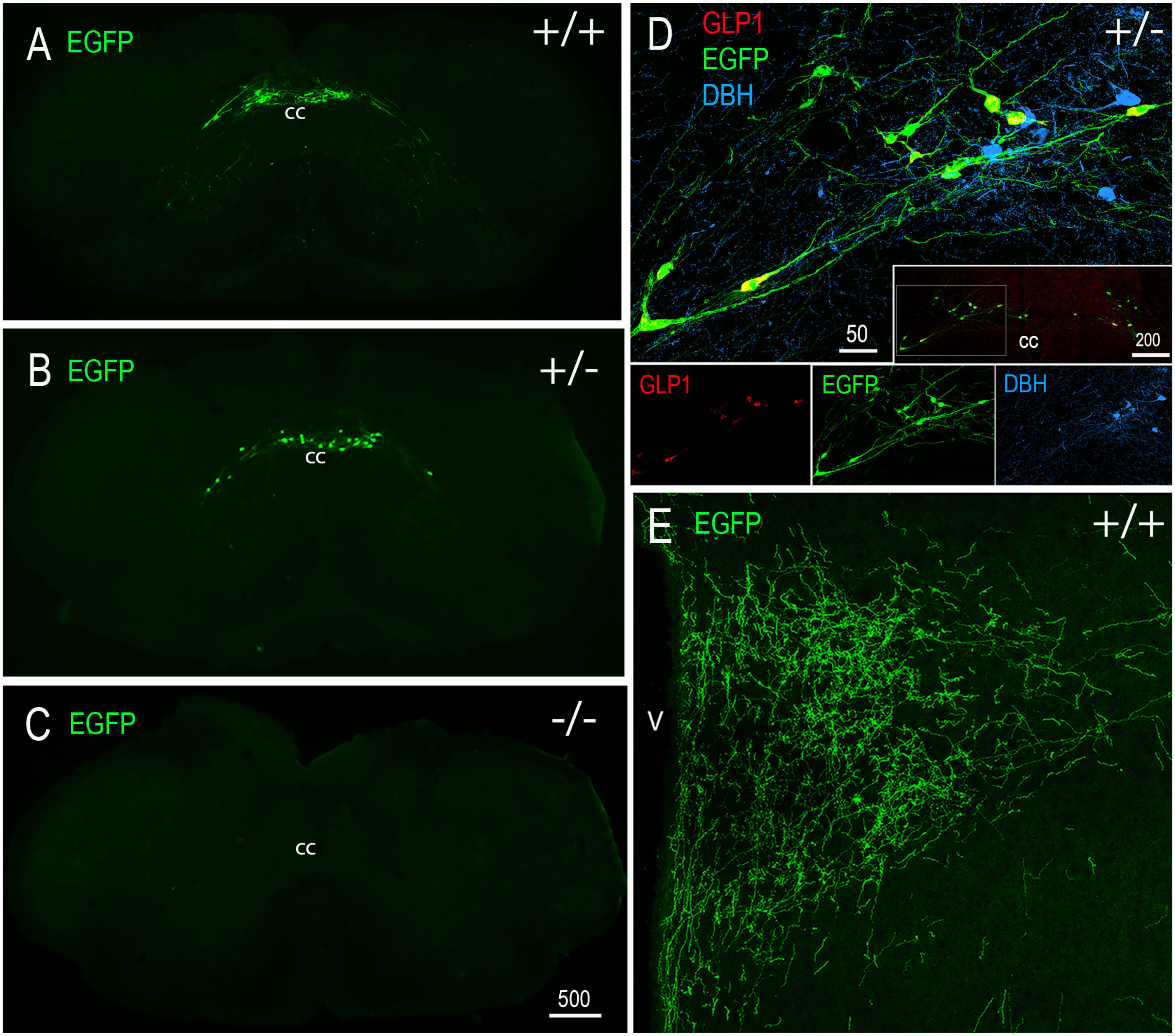

### Supp Fig. 4

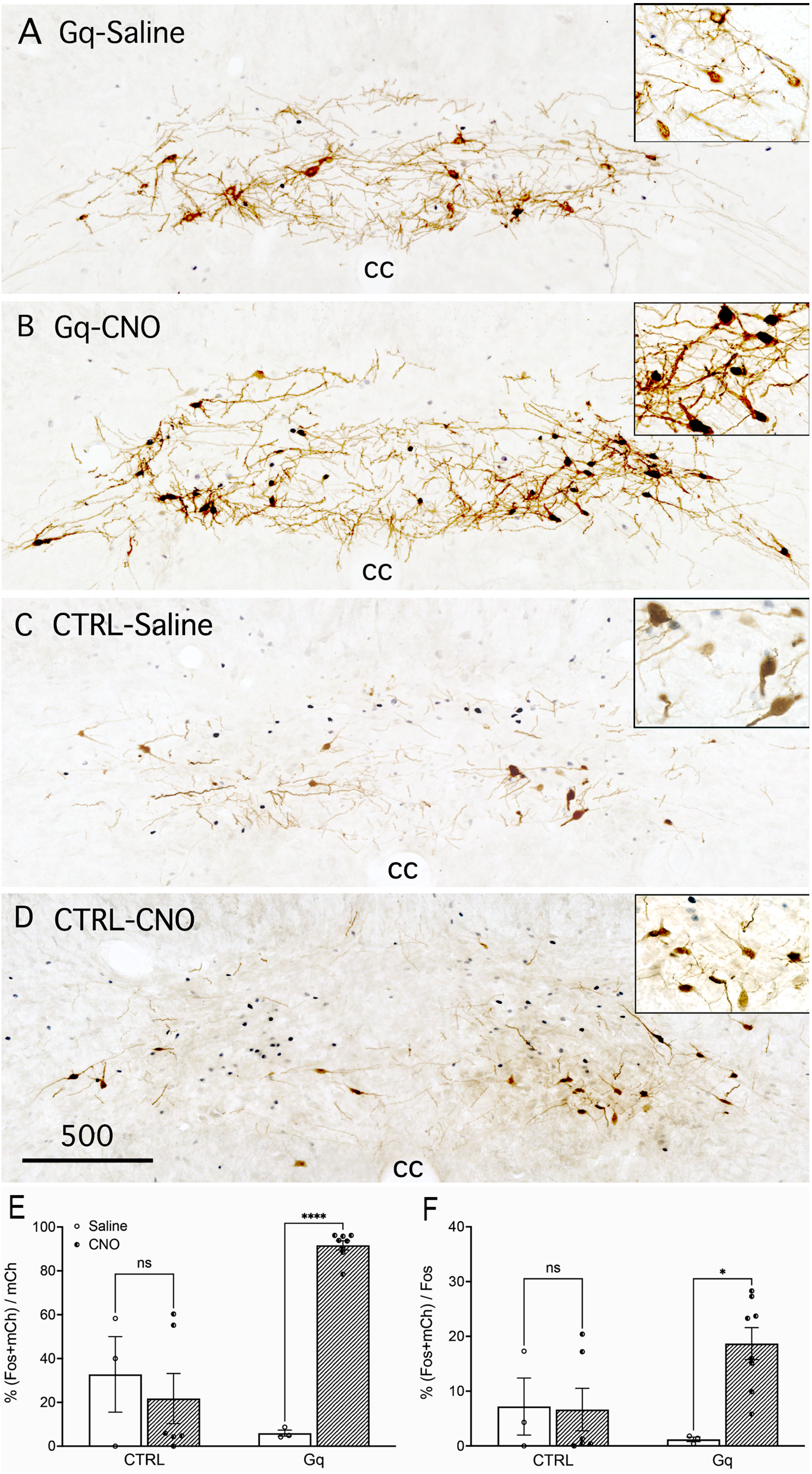

### Supp Fig. 5

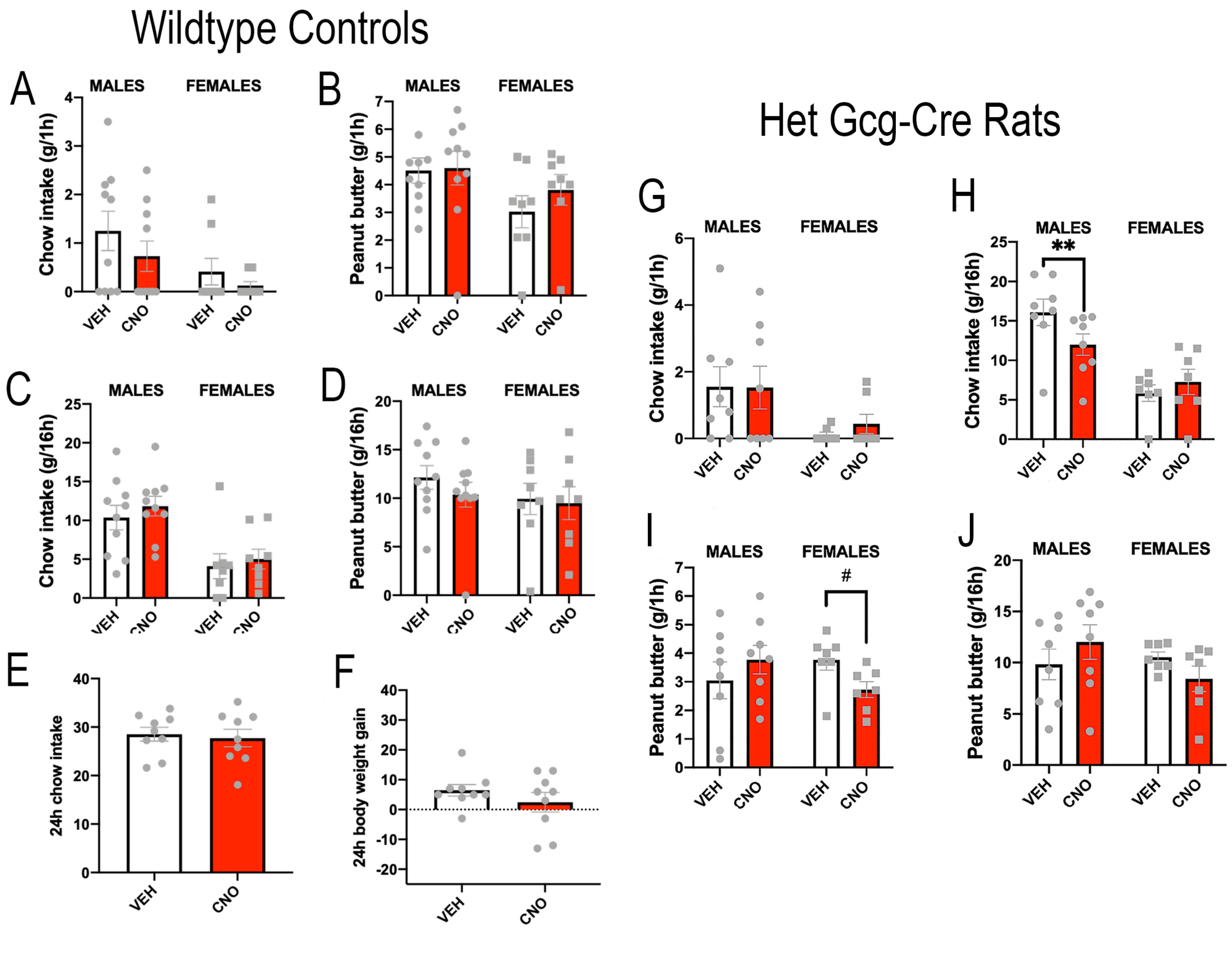
